# Ferroptosis propagates to neighboring cells via cell-cell contacts

**DOI:** 10.1101/2023.03.24.534081

**Authors:** Bernhard F. Roeck, Michael R. H. Vorndran, Ana J. Garcia-Saez

## Abstract

Ferroptosis is an iron-dependent form of regulated cell death characterized by accumulation of peroxidized lipids and plasma membrane disruption, whose molecular mechanism of execution remains poorly understood. Here, we developed a new optogenetic system, Opto-GPX4Deg, for light-induced degradation of the lipid reducing protein GPX4, which allows controlled ferroptosis induction with high precision in time and space. By using Opto-GPX4Deg to study cell death dynamics within the cellular population, we found that lipid peroxidation, followed by ferroptotic death, spread to neighboring cells in a distance-dependent manner. Remarkably, ferroptosis propagation showed a strong dependency on cell confluence and preferentially affected adjacent cells. Our findings establish cell death propagation as a feature of ferroptosis and provide new understanding of the mechanism involved.

## Introduction

Ferroptosis is an iron-depended form of regulated cell death that is mainly driven by the accumulation of phospholipid peroxides in cellular membranes^1,2^. However, ferroptosis is distinct from other regulated cell death pathways in that it is triggered when the redox defense systems in the cell are overruled by exceeding their antioxidant-buffering capabilities^3–5^. Thus, unlike apoptosis, pyroptosis or necroptosis, ferroptotic cell death lacks a terminal executioner protein^6^. While lipid peroxides accumulation has been identified as key event during ferroptosis, the molecular mechanisms governing ferroptosis regulation and execution remain poorly understood^7,8^. Ferroptosis has been associated with a number of diseases, including neurodegeneration, kidney failure or stroke^9–13^. In addition, induction of ferroptosis holds potential as an anti-cancer treatment strategy for refractory cancers. On the one hand, ferroptosis was shown to hinder cancer development through tumor suppressor engagement. On the other hand, some cancers are especially sensitive to ferroptosis due to their metabolic properties, their high ROS levels and specific mutations^14–16^. Elucidating the cellular mechanisms that regulate ferroptosis is hence of great relevance to devise new strategies for fighting human disease.

Four main ferroptosis defense mechanisms have been identified so far, which all have in common that they directly neutralize lipid peroxides, including the GPX4-GSH-, the FSP1-CoQH-, the DHODH-CoQH_2_ and the GCH1-BH_4_ systems^17^. Among these, the most effective strategy for inducing ferroptosis in therapy-resistant tumors is the inhibition of the enzyme GPX4, which disables the GPX4-GSH system – the most powerful anti-ferroptosis defense system^15^. GPX4 is a member of the GPX protein family, whose important function is to convert phospholipid hydroperoxides to phospholipid alcohols. This crucial role is emphasized by the fact that GPX4 depletion or inhibition triggers ferroptosis^18,19^. GPX4, together with its cofactor reduced glutathione (GSH), constitute the above mentioned GPX4-GSH system. GSH is a tripeptide consisting of glycin, glutamate and cysteine. Cysteine is the bottle neck in GSH synthesis and is mainly imported through the xc^-^ system as cystine, the oxidized form of cysteine^20^, which is then reduced to cysteine in the cytosol^21^. Disrupting either the cystine import through pharmacological inhibition of the xc^-^ system, e.g. with erastin, or depletion of cystine from the cell culture media was shown to induce ferroptosis in various cancer cell lines^1,22^.

To add another layer of complexity, ferroptosis seems to propagate through cell populations in vitro and has been linked with necrosis spread in diseased tissues^23–25^. Iron and lipid peroxidation have been shown to be required for ferroptosis propagation in cultured cells^26^. However, the use of drugs in previous studies for inducing ferroptosis, potentially affecting all cells in the population, has limited the investigation of ferroptosis spread, which remains under debate. Given the implications of this form of cell death in disease^23,25^, understanding the yet obscure mechanism underlying ferroptosis propagation is of key interest.

Here, we developed a new tool capable of inducing ferroptosis at will in selected cells by combining the so far most potent strategy for inducing ferroptosis, GPX4 depletion, with the advantages of optogenetics^15^. We designed an optogenetics construct, which we called Opto-GPX4Deg, that is based on light-induced proteasomal degradation of GPX4 leading to ferroptosis. The key advantages of Opto-GPX4Deg are unprecedented spatial and temporal control of ferroptosis induction, as well as the reversibility of the system compared to conventional drug treatments. Using this system, we show that light-controlled induction of ferroptosis occurred proportionally to the intensity and duration of the activating illumination, which could be rescued through the antioxidant ferrostatin in a concentration dependent manner. We prove the specificity of ferroptosis induction through C11-Bodipy staining, which is an established marker for lipid peroxidation, as well as by lipidomics analysis. Strikingly, we were able to show for the first time in an unbiased approach that ferroptotic cells are capable of inducing lipid peroxidation and ferroptosis in neighboring cells, which they further propagate to their adjoining cells.

## Results

### Opto-GPX4Deg induces cell death associated with ubiquitin mediated proteasomal degradation of GPX4

To devise an optogenetic system capable of inducing ferroptosis triggered by light, we designed a fusion construct aimed at depleting GPX4, the most potent protection protein against ferroptosis, in a light-controlled manner. To this aim, we generated an optogenetics tool based on a LOV2 domain that changes its conformation upon blue light illumination, making a RRRG degron sequence accessible for ubiquitin mediated proteasomal degradation, LOVpepdegron (**Fig. 1A**)^27,28^. The construct, which we called Opto-GPX4Deg, contained the cDNA of GPX4 tagged at the N-terminus with GFP for easy visualization of expression, and the LOVpepdegron at the C-terminus. As a negative control, we also generated a similar construct lacking the degron sequence, GFP-GPX4-LOVpep (named Opto-Ctrl).

**Figure 1:**
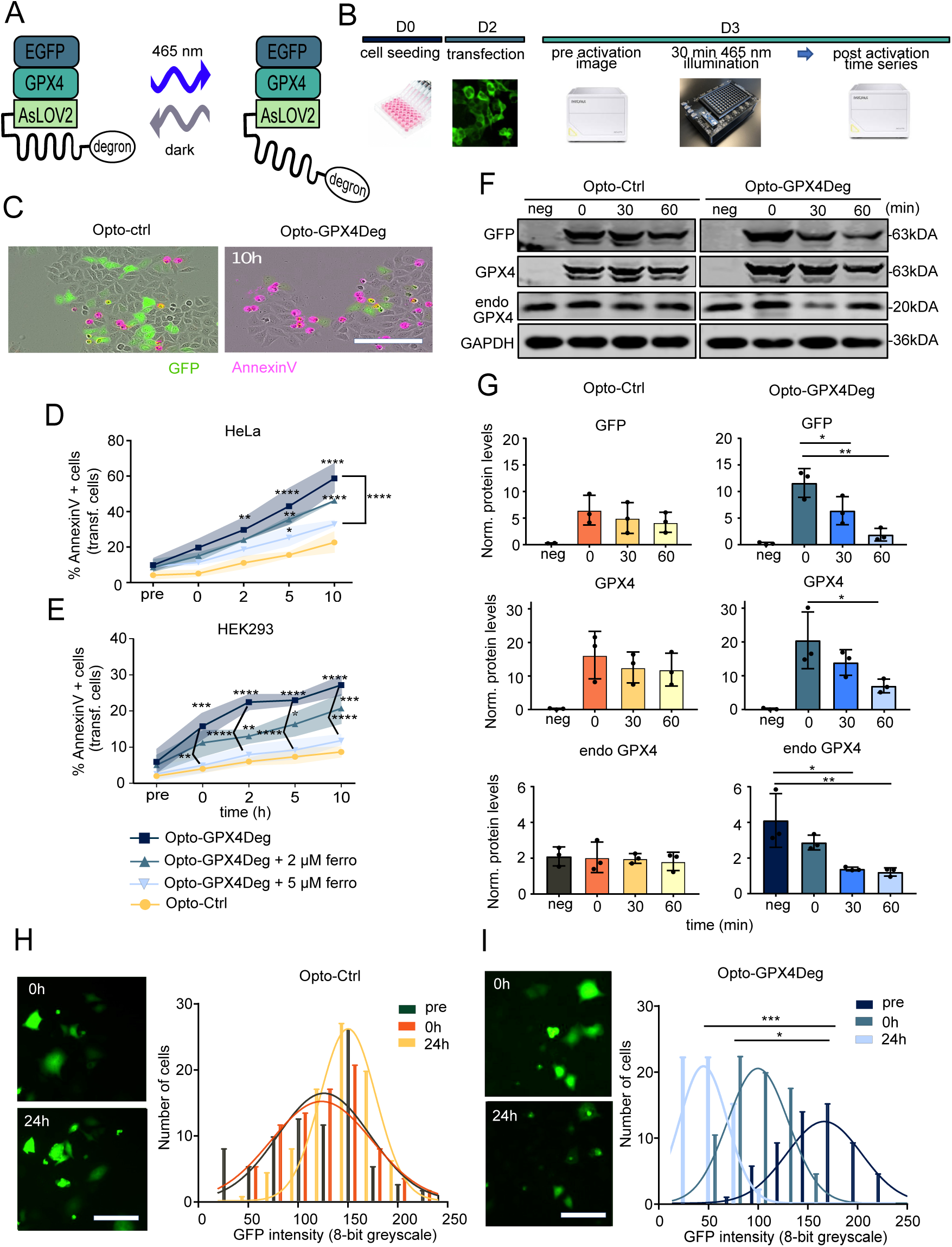
Ferroptosis induction via optogenetics. **(A)** Scheme of the Opto-GPX4Deg construct: The construct encodes for a fusion protein including EGFP, GPX4 and a LOV domain with a degron sequence on the C-terminal end. In the dark the degron is inaccessible. Upon activation with blue light the LOV domain changes its conformation making the degron sequence accessible for ubiquitin mediated proteasomal degradation.^27,28^ **(B)** Workflow of high-throughput optogenetics cell death experiments: On day 0, 1500 HeLa or HEK293 cells/well were seeded into a 96-well plate. On day 2, HeLa or HEK293 cells were transfected with Opto-Ctrl or Opto-GPX4Deg and incubated for 16h at 37°C. On day 3, the cells were either treated with 0, 2 or 5 µM of ferrostatin for 1h and 1:200 AnnexinV was added to the cells. Then a pre-activation image was acquired using a IncuCyte S3, followed by 30 min illumination with 100% power of 465 nm LEDs using an optoPlate-96.^29,30^ Subsequently, the 96-well plate was transferred to the IncuCyte S3 for acquiring a post-activation time series. **(C)** Representative merge images of 10h post-activation of HeLa cells transfected either Opto-GPX4Deg or Opto-Ctrl (green). Cell death was assessed by AnnexinV staining (magenta). Quantification of high-throughput optogenetics cell death experiment in HeLa **(D)** and HEK293 **(E)** cells. Experiments were performed as described in **(B)**. Cell death was assessed by AnnexinV staining and for the quantification of % AnnexinV positive cells, data were normalized to GFP positive cells. Statistical analysis by three-way ANOVA corrected for multiple comparisons using Tukey’s multiple comparison test. Asterisks indicate significant differences: *p < 0.05, **p < 0.01, ***p < 0.001, ****p<0.0001. n = 3 for all experiments. **(F)** Representative Western blotting showing the illumination dependent degradation of Opto-GPX4Deg: HeLa cells were either transfected with Opto-GPX4Deg or the Opto-Ctrl. After two days, the optogenetic constructs were activated either for 0, 30 or 60 min with 100% 465 nm LED intensity using an optoPlate-96. GFP as well as a GPX4 antibodies were used. GAPDH was used as loading control. **(G)** Quantification of the GFP, GPX4 and endogenous GPX4 proteins levels of the WB in **(F).** Protein levels were normalized to the respective loading control (GAPDH). Statistical analysis by one-way ANOVA corrected for multiple comparisons using Tukey’s multiple comparison test. Asterisks indicate significant differences: *p < 0.05, **p < 0.01, ***p < 0.001, ****p<0.0001. n = 3 for all experiments. Quantification of the GFP intensities for the experiments shown in **(C)** as a readout of the time dependent degradation of Opto-GPX4Deg. Statistical analysis by one-way ANOVA corrected for multiple comparisons using Tukey’s multiple comparison test. Asterisks indicate significant differences: *p < 0.05, **p < 0.01, ***p < 0.001, ****p<0.0001. n = 3 for all experiments. Values are displayed as mean ± SD.

We tested the functionality of the optogenetic tool, using a high-throughput optogenetics experiment pipeline in 96-well format. After confirming their sensitivity to ferroptosis **(Fig. S1)**, HEK293 or HeLa cells were transfected either with Opto-GPX4Deg or with Opto-Ctrl and cultured for 16h before being treated or not with different concentrations of ferrostatin. Then, we triggered the activation of the optogenetics constructs by illumination for 30 min using an optoPlate-96^29,30^(see Methods), and monitored the kinetics of cell death in the population via live cell imaging. As expected, HEK293 and HeLa cells expressing Opto-GPX4Deg died substantially more compared to cells expressing the Opto-Ctrl construct. Cell death induction upon Opto-GPX4Deg activation was proportional to the illumination intensity, which did not occur in cells expressing the Opto-Ctrl construct **(Fig. S2 A and B)**. Cell death induced by Opto-GPX4Deg could also be inhibited by ferrostatin treatment in a dose dependent manner (**Fig. 1C-E**), in support of ferroptotic cell death.

Although maintaining the anti-ferroptosis function of the Opto-GPX4Deg construct may not be necessary for its ability to induce ferroptosis, we confirmed that the fusion of the GFP and LOVpepdegron tags maintained, at least partially, the activity of GPX4. To this aim, we confirmed the ability of Opto-GPX4Deg expression to rescue cell death induced by either ferrostatin deprivation in GPX4 KO cells or by tamoxifen-induced GPX4 depletion **(Fig. S3 A-E)**, showing that the fusion protein retains partial activity.

We next validated that activation of the Opto-GPX4Deg tool indeed leads to the degradation of the fusion protein by Western Blot (WB). HeLa cells were transfected with the Opto-GPX4Deg or the Opto-Ctrl plasmid and then activated with different illumination durations. As shown by the signal reduction of GFP and GPX4 antibodies, the protein levels of Opto-GPX4Deg, but not the control construct, were significantly reduced (**Fig. 1F and G**). Interestingly, this trend could also be observed for the endogenous GPX4 levels, suggesting the existence of a coordinated mechanisms for the regulation of proteasomal degradation of GPX4. As additional proof of the light-induced degradation of the construct, we quantified the reduction in GFP intensity in individual cells expressing the Opto-GPX4Deg, but not the control construct, before, directly after and 24h post activation (**Fig1. H and I**, corresponds to experiments in **Fig. 1C**). Together, these results indicate that the Opto-GPX4Deg tool induces cell death by light-induced degradation of both exogenous and endogenous GPX4.

### Optogenetic ferroptosis induction can render cancer cell lines more sensitive to ferroptosis

The Opto-GPX4Deg system caused relatively more cell death in HeLa cells compared to HEK293 cells, which are more sensitive to pharmacological inhibition of GPX4 (**Fig.1** **C,D and S1**), suggesting the ability of the construct expression to affect ferroptosis sensitivity. We wondered whether other cancer cell lines would also be rendered more sensitive to ferroptosis by expression of Opto-GPX4Deg. To test this hypothesis, we first assessed the pharmacological ferroptosis sensitivity of three additional, different cancer cell lines (**Fig.2**). Small cell lung cancer (SCLC) cells turned out to be sensitive to ferroptosis induction only when high concentrations of RSL3 were used, whereas CT26 and B16-F10 cells were insensitive. Remarkably, we found that CT26 and B16-F10 cells transiently expressing the Opto-GPX4Deg construct underwent massive cell death upon illumination, which could again be delayed by the application of ferrostatin (**Fig.2** **A-L).** Most likely, this sensitizing effect could be caused by a downregulation of other cellular redox protection systems upon transient overexpression of Opto-GPX4Deg, combined with degradation of the construct upon optogenetic activation and the simultaneous downregulation of endogenous GPX4 **(Fig. 1F-I and Fig. S3**). Furthermore, these experiments also show that Opto-GPX4Deg can induce light-controlled cell death in a cell type-independent manner.

**Figure 2:**
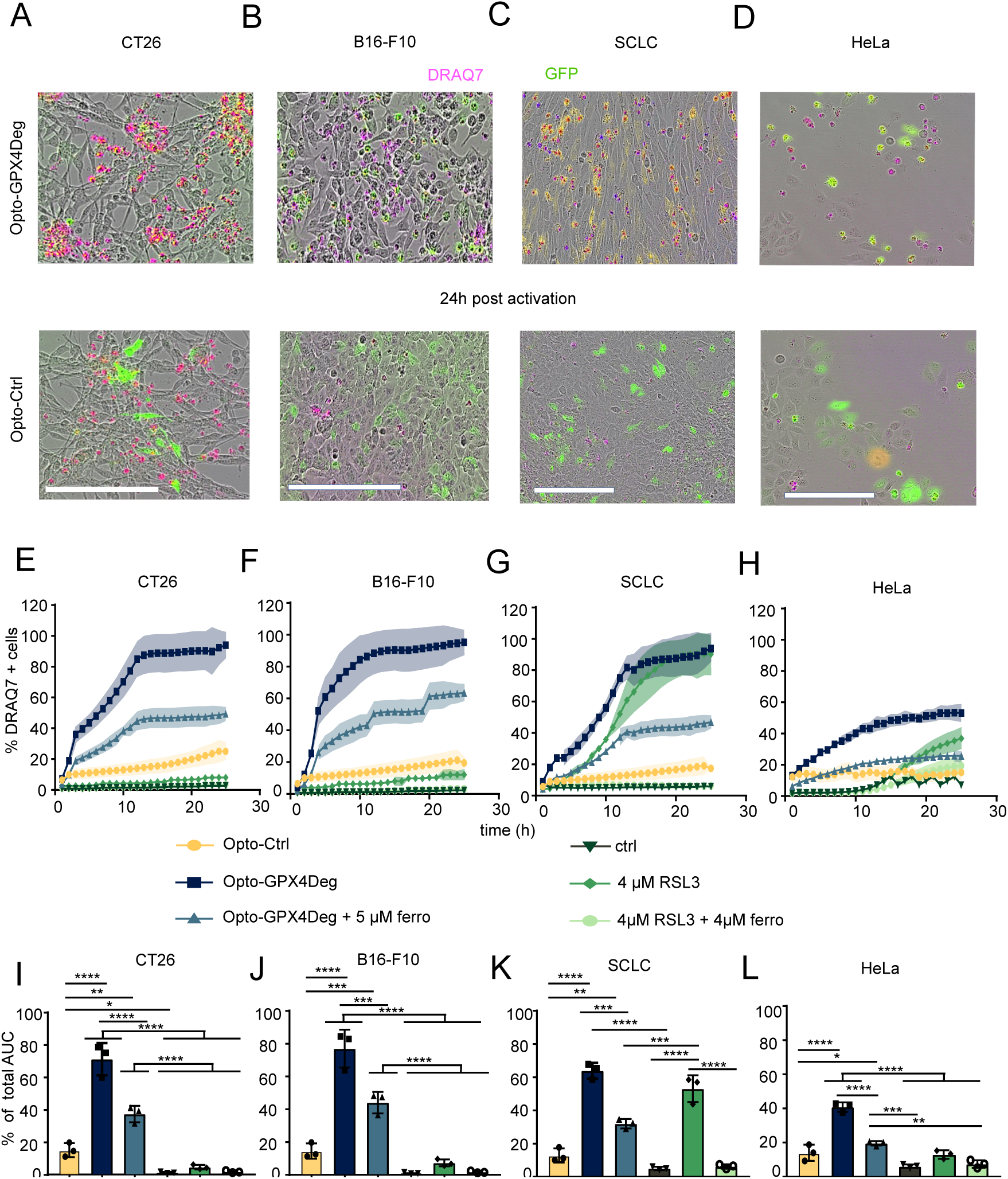
Opto-GPX4Deg enables increased efficacy in killing various cancer cell lines compared to pharmacological GPX4 inhibition. On day 0, 10k CT26, SCLC, B10-F10 or HeLa cells per well were seeded into a 96-well plate. On the next day the cell lines were treated with different combination treatments: RSL3, RSL3 + ferrostatin or DMSO in the in indicated concentrations. 1:200 Draq7 was added for cell death assessment. Optogenetic experiments were performed as described in Fig. 1B with the difference that here 10.000 CT26, SCLC, B10-F10 or HeLa cells were seeded into 96-well plate and subsequently incubated for 2 days before being transfected with Opto-GPX4Deg or Opto-Ctrl for 16h. 1,5% 465 nm LED intensity for the CT26, B16_F10 and SCLC was used for activation. On the next day, the cell lines were treated or not with 5 µM ferrostatin. 1:200 Draq7 was added for cell death assessment using a IncuCyte S3. **(A-D)** Representative merge images of 24h post-activation of CT26, B16-F10, SCLC or HeLa cells transfected either with Opto-GPX4Deg or the Opto-Ctrl (green). Cell death was assessed by Draq7 staining (magenta). Scale bars, 200 µm. **(E-F)** Quantification of the cell death kinetics. (**I-L**) To assess statistical differences, the % total area under the curve of the different treatments was calculated and subsequently a one-way ANOVA was performed and corrected for multiple comparisons using Tukey’s multiple comparison test. Asterisks indicate significant differences: *p < 0.05, **p < 0.01, ***p < 0.001, ****p<0.0001. n = 3 for all experiments. Values are displayed as mean ± SD.

### Opto-GPX4Deg allows monitoring ferroptosis with high spatial and temporal resolution

One advantage of optogenetic tools is that they can be combined with live cell microscopy to investigate the consequences of activating signaling pathways with high temporal and spatial resolution. We thus optimized the experimental procedure to image the dynamics of ferroptotic cell death induced with Opto-GPX4Deg at the single cell level using live cell confocal microscopy (**Fig. 3A**). We acquired a pre-activation image followed by Opto-GPX4Deg activation using the FRAP unit of the microscope with 100 iterations of illumination at 70% intensity of a 405 nm laser every 5 min. As expected, activating illumination of selected, individual HEK293 cells expressing Opto-GPX4Deg efficiently induced cell death, which could be detected with unprecedented spatial and temporal precision (**Fig. 3B**).

**Figure 3:**
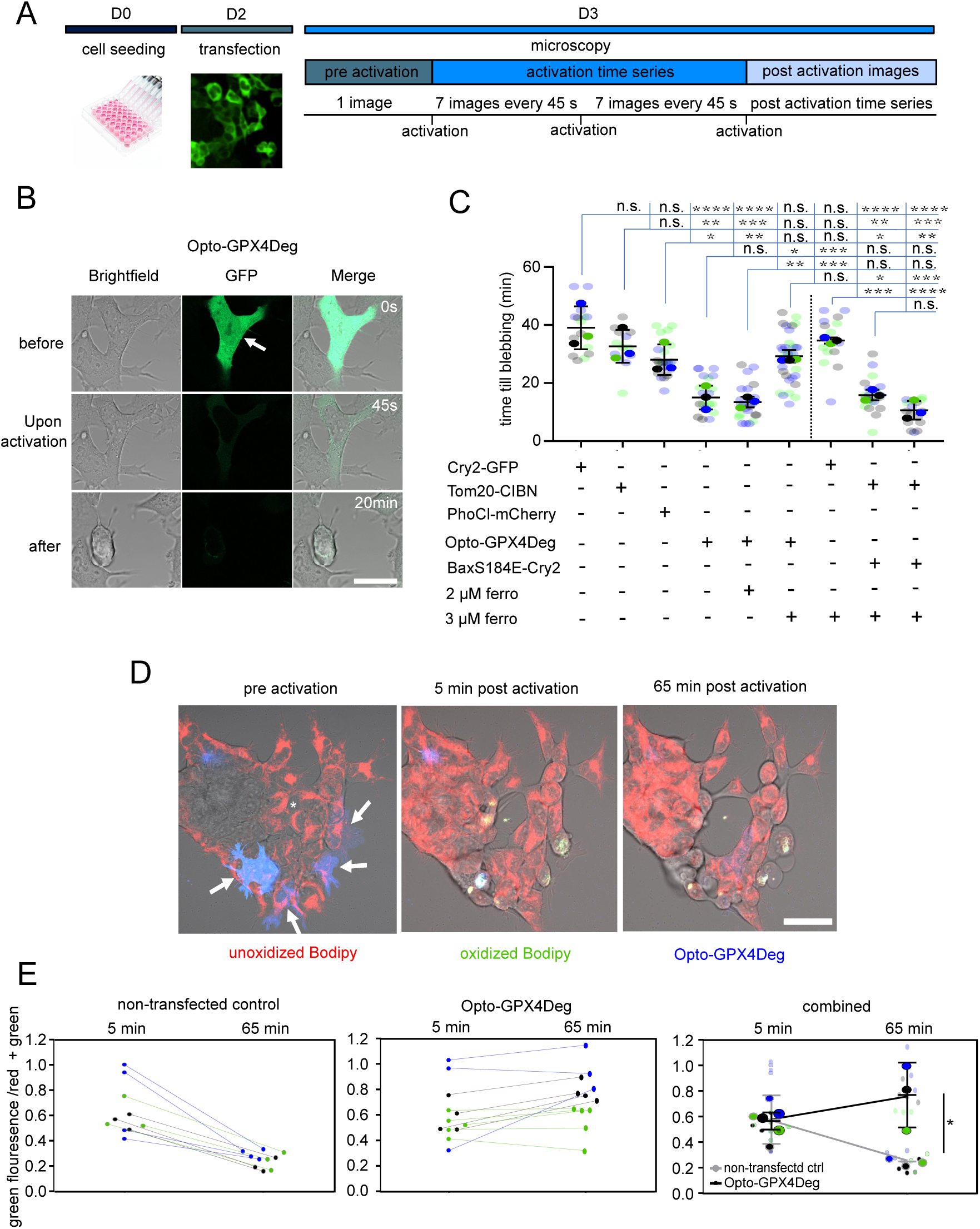
Single cell ferroptosis induction with Opto-GPX4Deg. **(A)** Workflow of confocal optogenetics cell death experiments: On day 0, 5.000 HEK cells were seeded into a 8-well chamber slide. On day 2, HEK293 cells were transfected with different optogenetic tools and incubated for 16h at 37°C. On day 3, the cells were either treated with 0, 2 or 3 µM ferrostatin for 1h and brought to a confocal microscope for optogenetic activation. Before activation, one pre-activation image was acquired followed by the optogenetic activation using the FRAP unit with 70% intensity of a 405 nm laser with 100 iterations every 5 min. After optogenetic activation, images were acquired every 45s. **(B)** Representative images showing two cells expressing Opto-GPX4Deg upon activation (white arrow). Scale bars, 10 µm. **(C)** Quantification of the time till blebbing as proxy for cell death: HEK293 cells were transfected with the indicated constructs (first three lanes represent different optogenetic control constructs), treated or not treated with different concentrations of ferrostatin and activated as described in **(A).** To test the influence of bleaching-induced ROS production Cry2_GFP (control) and an optogenetic Bax system were included in the analysis (last three lanes) and treated or not with 3 µM ferrostatin. Big dots represent the mean of three independent experiments, with small dots representing single cells of one individual experiment. Error bar = SD. **(D-E)** Optogenetic induction of Opto-GPX4Deg results in increased levels of lipid peroxidation. For normalization, the green (oxidized) Bodipy fluorescence signal was divided by the green + red Bodipy fluorescence signal. White asterisk indicates activated, non-transected controls and white arrows indicate activated cells expressing Opto-GPX4Deg. Scale bar, 50 µm. The big 3 dots represent the mean of three independent experiment and the small data dots represent single cells of one individual experiment. For assessing statistical differences a one-way ANOVA was performed and corrected for multiple comparisons using Tukey’s multiple comparison test. Asterisks indicate significant differences: *p < 0.05, **p < 0.01, ***p < 0.001, ****p<0.0001. n = 3 for all experiment. Values are displayed as mean ± SD.

To quantify cell death induction at the single cell level, we measured the time lag between the activating illumination and plasma membrane blebbing in the illuminated cells, which spanned a narrow distribution of around 13-20 minutes of illumination, with an average of 17 minutes (**Fig. 3C**). When compared to cells expressing negative controls, cell death induction occurred significantly faster in activated cells expressing Opto-GPX4Deg, indicating specific induction of cell death. The negative controls also provided an estimation of the photo-toxicity by the activating illumination. This is an important parameter to consider when optimizing illumination conditions in order to avoid any overlap between optogenetics-induced cell death and photo-toxicity.

In agreement with the high throughput assay (**Fig.1**), treatment with 3 µM ferrostatin was capable of rescuing cell death induction to control levels, again supporting that activated cells expressing Opto-GPX4Deg die by ferroptosis. Yet, ferrostatin treatment did not affect the time till blebbing due to photo-toxicity in control samples, nor cell death induction using an already established apoptosis optogenetics tool.^31^

### Light-induced activation of Opto-GPX4Deg promotes lipid peroxidation – a hallmark of ferroptosis

To validate that light-controlled activation of Opto-GPX4Deg induces cell death by ferroptosis, we measured the induction lipid peroxidation in cellular membranes, a hallmark of this form of cell death. We transfected HEK293 cells with Opto-GPX4Deg and stained them with C11-Bodipy, a membrane-bound ratiometric fluorescent indicator of lipid oxidation, commonly used as a proxy for ferroptosis induction. We calculated the ratio between the green fluorescence emission (oxidized form of the dye) and the total emission in the green and red channels (oxidized plus reduced forms) at 5 min and 65 min after activating illumination of the individual cells.

Interestingly, the activating illumination caused a wave of C11-Bodipy oxidation both in the non-transfected controls and in the cells expressing Opto-GPX4Deg, possibly as a consequence of the exposure to elevated light intensity. However, the C11-Bodipy oxidation ratio was reduced to normal levels after 1h in the non-transfected cells, whereas cells expressing Opto-GPX4Deg maintained or slightly increased the oxidation ratio (**Fig. 3D-E**). Moreover, the non-transfected cells did not change their morphology over time, while the cells expressing Opto-GPX4Deg showed typical morphological features of ferroptosis, such as plasma membrane ballooning (**Fig.3** **D)**. Similar results were obtained when we measured lipid peroxidation with C11-Bodipy in the high throughput assay with the optoPlate-96 **(Fig. S4**).

We further validated that cell death induced by Opto-GPX4Deg was ferroptotic by lipidomic analysis. To this aim, we upscaled the format of the high-throughput experiment in **Fig. 1B**. Cells were incubated for two days in the dark before activating or not the Opto-GPX4Deg tool or Opto-Ctrl. Two days later the cells were harvested and processed (**Fig. 4A**). WB analysis confirmed the degradation of endogenous GPX4 and Opto-GPX4Deg, but not the Opto-ctrl construct (**Fig. 4B and C**). Importantly, the mass spectrometry analysis revealed a significant increase in oxidized lipid species in the Opto-GPX4Deg samples, but not in the negative controls (**Fig. 4D**). We detected the highest fold enrichments in oxidized phosphatidylcholine (PC) species, in line with previous work in HeLa cells.^32^ These observations were accompanied by a global reduction in fatty acids species (**Fig. 4E**), also in good agreement with lipidomics analysis of ferroptotic HT-1080 cells, where numerous mono-unsaturated fatty acids were depleted.^33,34^ Together, these data indicate that cell death induction by Opto-GPX4Deg is accompanied with elevated levels of lipid peroxidation compared to the controls, providing further evidence for ferroptosis induction.

**Figure 4:**
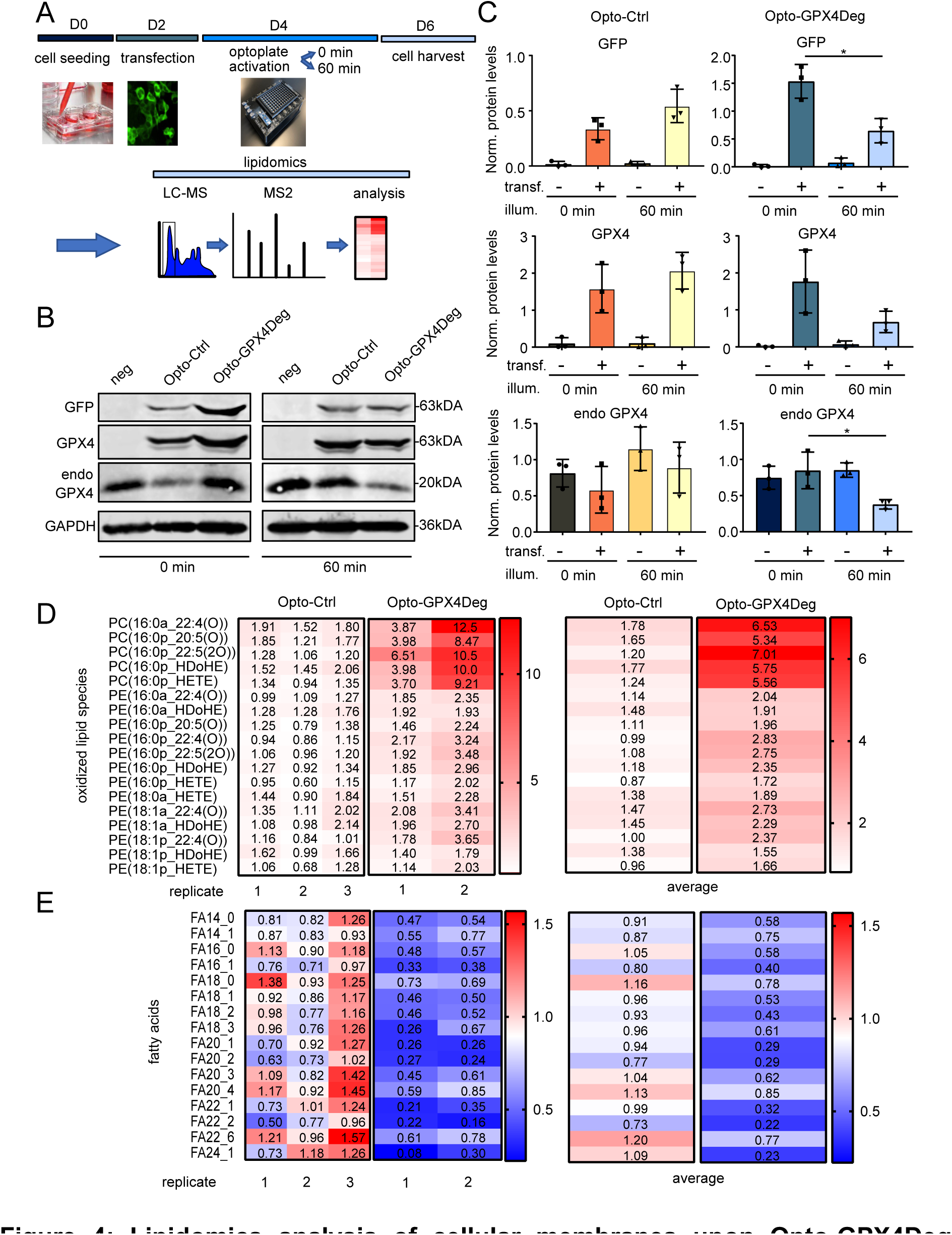
Lipidomics analysis of cellular membranes upon Opto-GPX4Deg activation. **(A)** Workflow of lipidomic experiments: On day 0, 100.000 HeLa cells/well were seeded into 6-well plates. On day, 2 HeLa cells were transfected either with Opto-Ctrl, Opto-GPX4Deg or not transfected and incubated for 48h at 37°C. Then the bulks were illuminated or not for 60 min with 100% 465 nm LED intensity using an optoPlate-96. After 48h, the cells were harvested and samples were prepared for WB and lipidomics. **(B)** Representative WB showing the illumination-dependent degradation of the Opto-GPX4Deg construct. GFP as well as GPX4 antibodies were used and GAPDH was used as loading control. **(C)** Quantification of the GFP, GPX4 and endogenous GPX4 proteins levels of the WB in **(B).** Protein levels were normalized to the respective loading control (GAPDH). –, non-transfected, +, transfected either with Opto-Ctrl or Opto-GPX4Deg. Statistical analysis by one-way ANOVA corrected for multiple comparisons using Tukey’s multiple comparison test. Asterisks indicate significant differences: *p < 0.05, **p < 0.01, ***p < 0.001, ****p<0.0001. n = 3 for all experiments. Values are displayed as mean ± SD. **(D)** Analysis of oxidized lipids was performed according to^4^. Fold change in the heat map calculated by dividing Area ratios/µg protein of different oxidized lipid species upon illumination divided by the non-illuminated control for the individual replicates (left) and the average (right). **(E)** Fatty acids analysis was performed according to^4^. Fold change in the heat map calculated by dividing Area ratios/µg protein of different oxidized lipid species upon illumination divided by the non-illuminated control for the individual replicates (left) and the average (right).

### Ferroptotic cells have the capacity to induce cell death in neighboring cells

Previous reports have proposed that ferroptosis can propagate within the cell population in vitro and that it contributes to necrosis propagation in diseased tissues via yet unknown mechanisms^23,26^. However, this notion is not fully established because conventional ferroptosis induction strategies with drugs affect all cells in the population, allowing only restricted conclusions about the responses of neighboring cells. We reasoned that our Opto-GPX4Deg system would enable studying ferroptosis propagation to neighboring cells without these limitations. To this aim, we quantified the cell death induction over time in the population of transfected and non-transfected HeLa cells upon Opto-GPX4Deg activation. Intriguingly, we found that not only cells expressing Opto-GPX4Deg, but also non-transfected cells, died upon illumination, which did not happen in the samples transfected with the Opto-Ctrl construct (**Fig. 5A - D**).

**Figure 5:**
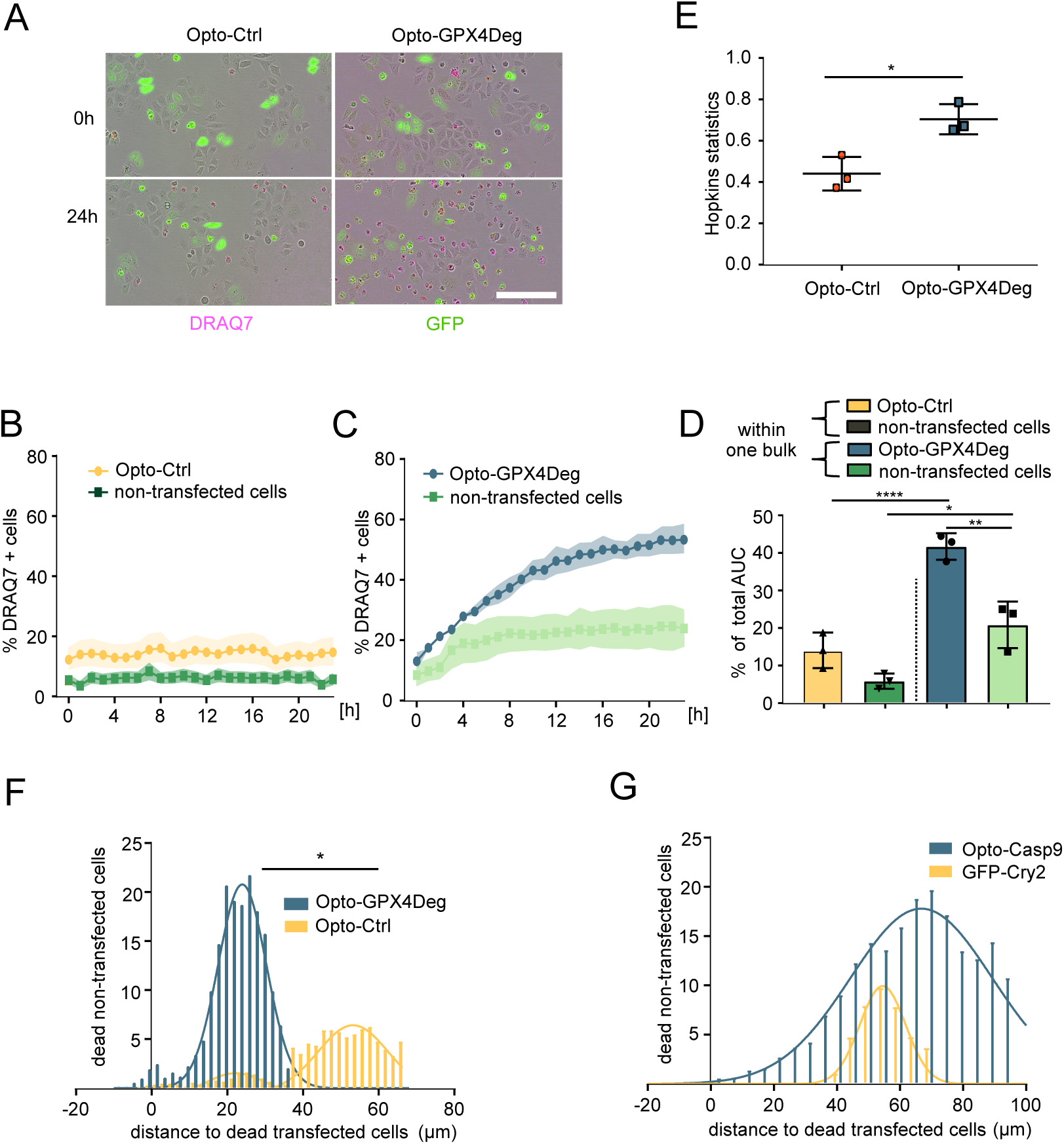
Ferroptotic cells induce cell death in neighboring cells. Experiments performed according to workflow in Fig. 1B, except that here a 12-well plate with a seeding density of 100.000 HeLa cells/well was used. Cell death was assessed by Draq7 staining. % Draq7 positive cells that were GFP negative, as well as GFP positive, calculated with custom made software. (**A**) Representative images of 0h and 24h post-activation showing increased cell death in non-transfected cells in the Opto-GPX4Deg transfected sample compared to the Opto-Ctrl sample. **(B-D)** For assessing statistical differences of cell death induction, the % total Area under the Curve of the different cell populations was calculated and subsequently a one-way ANOVA was performed and corrected for multiple comparisons using Tukey’s multiple comparison test. Asterisks indicate significant differences: *p < 0.05, **p < 0.01, ***p < 0.001, ****p<0.0001. n = 3 for all experiments. (**E**) Hopkins statistical analysis was performed to assess the cluster tendency of the datasets. To assess statistical differences a parametric t-test was performed. Asterisks indicate significant differences: *p < 0.05, **p < 0.01, ***p < 0.001, ****p<0.0001. n = 3 for all experiments. **(F)** Data derived from experiments in **(A-E).** Quantification of the distance distribution of either dead cells expressing Opto-GPX4Deg or dead cells expressing Opto-Ctrl to dead, non-transfected cells. Same procedure as in **(G)**, but here optogenetic constructs for inducing apoptosis and the respective controls were used. To assess statistical differences, a parametric t-test was performed. Asterisks indicate significant differences: *p < 0.05, **p < 0.01, ***p < 0.001, ****p<0.0001. n = 3 for all experiments. Values are displayed as mean ± SD.

Analysis of the cell death kinetics suggested that, upon activation, cells expressing the Opto-GPX4Deg tool died first, followed by the non-transfected cells (**Fig. 5A-C**). If the death of the non-transfected cells were unrelated to the activation of Opto-GPX4Deg, one would expect a homogenous distribution of cell death within the non-transfected cell population. To check whether cell death was randomly distributed in the population or segregated into clusters, we then calculated the clustering tendency of our dataset using Hopkins statistics. Hopkins statistics is a measure of data clustering based on the difference between the distance from a real point to its nearest neighbor compared to the distance from a randomly chosen point within the data set to the nearest real data point. If the data has little to no clustering, the distance from one real point to another is on average the distance from a uniformly distributed random point to a real point, meaning that the value of Hopkins statistic will be around to 0.5. If the data are organized in tight clusters the distance from a real data point to its nearest neighbor will be small compared distance from an artificial point to a real data point, giving a value close to 1.0^35^. Interestingly, Hopkins analysis revealed that the cells in the population of the Opto-GPX4Deg samples died preferentially in clusters (value =0,67), whereas cell death was randomly distributed (value = 0,43) in the population of cells transfected with the Opto-ctrl (**Fig. 5E**).

Following these results, we explored whether cells dead by Opto-GPX4Deg activation were inducing cell death preferentially in cells located in close proximity. We quantified the mean distance between dead cells expressing Opto-GPX4Deg and non-transfected dead cells. Strikingly, we found that the distance from the dead non-transfected cells to the dead transfected cells was significantly shorter in the samples expressing Opto-GPX4Deg compared to the control construct (**Fig. 5F**). Furthermore, the distance in the Opto-GPX4Deg samples was in good agreement with the average diameter of HeLa cells (that we estimated at around 23 µm for dead and living cells combined, **Fig. S5**), suggesting that cells dying by ferroptosis induced by Opto-GPX4Deg were capable of inducing cell death in bystander cells in the immediate vicinity. This behavior was not observed when inducing apoptosis via optogenetics (**Fig. 5G**), suggesting that the propagation of cell death to neighboring cells is a specific trait of ferroptosis.

To dissect whether the distance between cells is a relevant parameter for ferroptosis propagation, we seeded HeLa cells in different confluences, transfected them with Opto-GPX4Deg and subsequently activated them with the optoPlate-96. In support of this hypothesis, we found that increased cell confluence rendered non-transfected HeLa cells more sensitive to cell death induced by neighboring cells dying through Opto-GPX4Deg activation **(Fig. S6A)**. As additional control, we seeded HeLa cells in different confluences and treated them with RSL3, which reproduced the same trend of higher cell death sensitivity by higher cell confluence **(Fig. S6B).** However, at a confluence of 100%, HeLa cells became insensitive to ferroptosis induction by RSL3, which was also observed in the samples transfected with Opto-GPX4Deg. This is in agreement to reports that extremely high cell densities promote the survival of GPX4 knockout (KO) cells^36^. Taken together, these data demonstrate in an unbiased approach that ferroptotic cells have the capacity to induce cell death in neighboring cells and strongly suggest that cell death propagation takes places between cells in close proximity.

### Cells adjacent to ferroptotic cells die by ferroptosis and are capable of further propagating ferroptosis to their neighbors

We next investigated the form of cell death experienced by non-transfected cells neighboring cells dying via Opto-GPX4Deg activation. For this purpose, we measured the kinetics of C11-Bodipy oxidation and cell death in a high-throughput assay as in **Fig. S4**, but this time simultaneously assess the signals of reduced and oxidized C11-Bodipy, as well as the BFP2 signal of a version of the Opto-GPX4Deg construct in which GFP had been exchanged with BFP2. This experimental system allows dissecting the kinetics of membrane oxidation and cell death in transfected and non-transfected cells, as well as the quantification of distances between dead, transfected cells and dead, non-transfected cells using our custom-made software in a single experiment.

We found that cells expressing the activated Opto-GPX4Deg tool first accumulated oxidized C11-Bodipy, followed by plasma membrane bursting and cell death. With a delay, direct neighboring cells to these ferroptotic cells also accumulated first oxidized lipids and eventually died. Interestingly, this was followed by increased lipid oxidation and cell death in their adjoining cells (**Fig. 6A** **and Fig. S7**). These results indicate that non-transfected cells neighboring to cells dying by Opto-GPX4Deg activation also died by ferroptosis, and that they were capable of further propagating lipid oxidation and ferroptotic cell death to other non-transfected cells in their close proximity. This system recapitulated the cell death kinetics shown in **Fig. 5A-C**, with activated Opto-GPX4Deg expressing cells dying first, followed by the non-transfected neighboring cells with a delay of around 3,5 h. In the control samples (cells non-transfected or transfected with the control construct BFP2_LOVpepdegron), only minor levels of lipid peroxidation were detected and no cell death propagation was observed **(Fig. S8)**.

**Figure 6:**
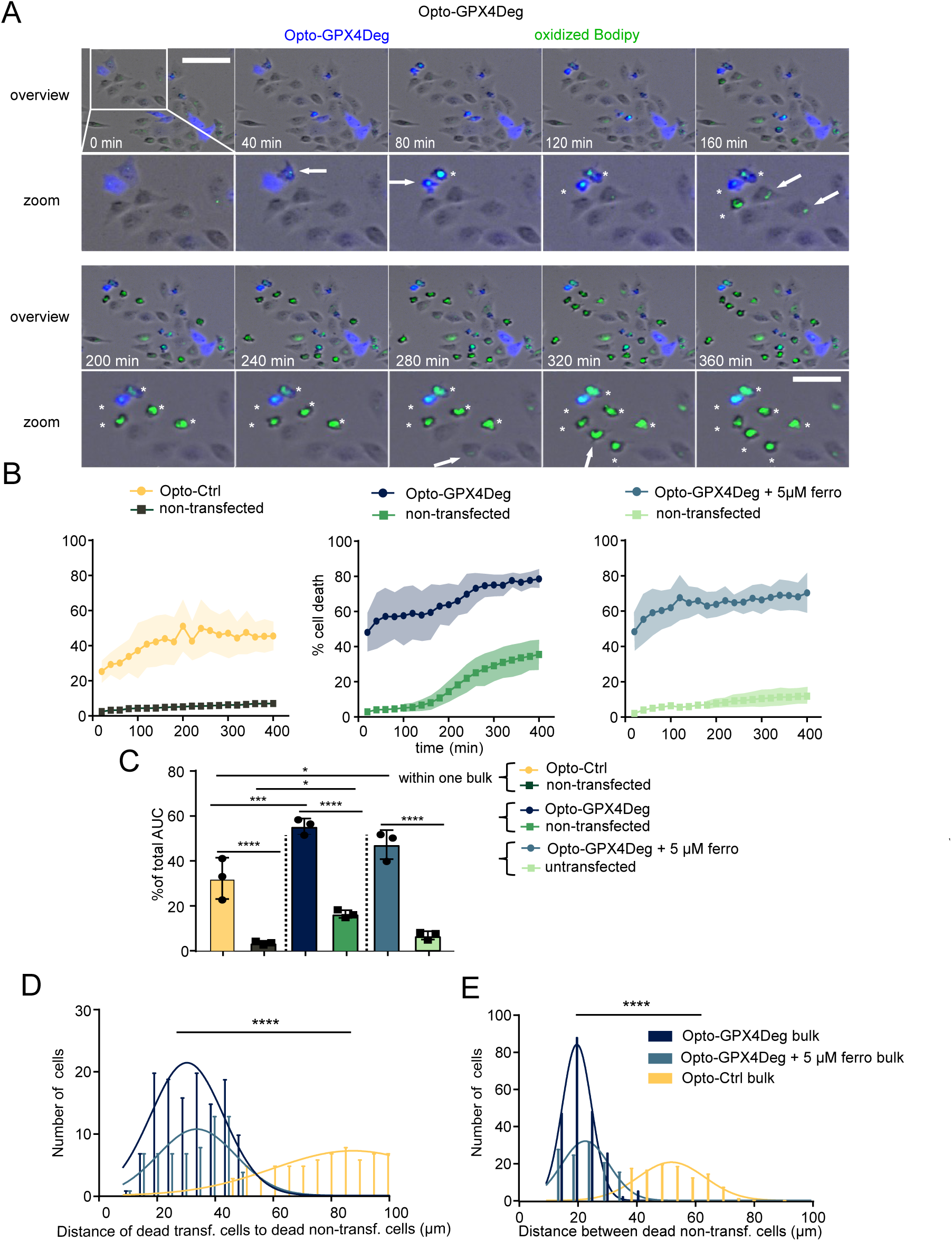
Cells adjacent to ferroptotic cells die by ferroptosis and are capable of propagating ferroptosis to their neighbors. Bodipy and cell death kinetics using the ImageXpress Micro 4: On the first day, 50.000 cells were seeded into 24-well plates, transfected or not on the next day with Opto-GPX4Deg or Opto-Ctrl and incubated for 16h at 37°C. On the next day, HeLa cells were stained with 1 µM Bodipy for 1h and then activated for 30 min with 100% 465 nm LED intensity using an opto-plate-96. Upon activation, cells were treated or not with 5 µM ferrostatin. **(A)** Representative time series of an activated sample expressing Opto-GPX4Deg. Blue signal, Opto-GPX4Deg. Green signal, oxidized Bodipy. White box indicates zoomed in area of interest (lower panels). Green arrow indicates lipid peroxide accumulation. White asterisks indicate dead cells. Scale bars, 100 µm for the overview and 50 µm for the zoomed in area. **(B and C)** Cell death quantification using custom made homemade software. For assessing statistical differences, the % total Area under the Curve of the different cell populations was calculated and subsequently a one-way ANOVA was performed and corrected for multiple comparisons using Tukey’s multiple comparison test. Asterisks indicate significant differences: *p < 0.05, **p < 0.01, ***p < 0.001, ****p<0.0001. n = 3 for all experiments. Values are displayed as mean ± SD. **(D)** Quantification of the distances of dead, transfected cells to dead, non-transfected cells. **(E)** Same as in **(D)**, but here the distances between dead, non-transfected cells was quantified.

We then administered ferrostatin to samples after Opto-GPX4Deg activation in order to test whether non-transfected neighboring cells were protected. Because cells were treated with ferrostatin only after optogenetic activation, the Opto-GPX4Deg expressing cells showed the same cell death kinetics as without ferrostatin treatment, whereas cell death of non-transfected cells was largely diminished, indicating that cells adjoining to dead, ferroptotic cells also die by ferroptosis (**Fig.6B-C**).

Next, we wanted to test whether these dying, adjoining cells are also capable of propagating ferroptosis to their neighboring cells. To this aim, we quantified the distance between dead, non-transfected cells in the Opto-GPX4Deg samples as well as in the different control samples. In line with our hypothesis, the average distance between dead cells corresponded to that of neighboring cells. This effect was again largely absent in the control samples either treated with ferrostatin or when transfected with the control construct (**Fig. 6E**). Together, these results show that cells adjoining ferroptotic cells die by ferroptosis and that they can propagate ferroptotic death to cells in their vicinity.

## Discussion

In this study, we developed, to our best knowledge, the first optogenetics tool capable of inducing ferroptosis, which we called Opto-GPX4Deg. It exploits the most potent strategy known to date for triggering this form of cell death, namely the depletion of GPX4, thus disrupting the GPX4-GSH redox defense system^15^. Opto-GPX4Deg is based on a fusion protein encoding for GFP, GPX4, as well as a LOV2 domain modified with a RRRG degron that becomes exposed and robustly induces ubiquitin mediated proteasomal degradation of the fusion construct when illuminated with blue light^27,28^.

We confirmed the light-induced degradation of the GPX4 fusion protein by WB and fluorescence analysis. Interestingly, we observed that the levels of endogenous GPX4 were also reduced upon illumination in cells expressing Opto-GPX4Deg, supporting a coordinated mechanism of regulation. In this scenario, cell death induction by Opto-GPX4Deg can be explained by the degradation upon light activation of the partly active GFP-GPX4-LOVpepdegron fusion protein, accompanied by downregulation of endogenous GPX4, leading to sufficient disruption of cellular redox systems for triggering ferroptosis.

We validated that Opto-GPX4Deg can be used for light-controlled induction of cell death exhibiting the key hallmarks of ferroptosis in different cell lines, both in single cell microscopy experiments as well as in high throughput assays of entire cell populations. Accordingly, light-induced cell death by Opto-GPX4Deg could be inhibited by ferrostatin in a dose dependent manner. In addition, it was associated with the appearance of peroxidized lipids in illuminated cells specifically expressing Opto-GPX4Deg, which we detected by both C11-Bodipy fluorescence as well as by lipidomics analysis.

The lipidomics analysis revealed significantly increased levels of various oxidized species of phosphatidylethanolamine (PE) and phosphatidylcholine (PC) in cells dying upon activation of Opto-Ferroptosis. PEs and PCs are the most abundant lipid species in plasma membranes, although there are differences in lipid composition between different cell types^37^. Currently, oxidation of phospholipids in ferroptosis is considered as a semi-specific process with some lipids being more prone to the oxidative process than others. Poly-unsaturated PEs have been identified as a crucial substrate for oxidation during ferroptosis, which is reflected in our data^38,39^. In our data oxidized forms of PCs were most upregulated, which is in agreement with ferroptosis work in HeLa cells.^32^ We also detected a general trend of fatty acids loss, including some poly-unsaturated free fatty acids. Similar results have been found in erastin-treated HT-1080 cells, where numerous mono-unsaturated fatty acids were depleted, suggesting that their peroxidation could occur downstream of initial poly-unsaturated lipid oxidation^33,34^. It will be interesting to investigate whether the specific substrates for lipid oxidation during ferroptosis are preferred depending on cell type and/or metabolic conditions.

Remarkably, we found that Opto-GPX4Deg expression could sensitize several cancer cell lines to light-induced ferroptosis, which opens new opportunities to explore the use of this strategy for anti-cancer treatments. In combination with delivery methods into tumor cells, such as those used in RNA vaccines^40^, the high spatial and temporal control of Opto-GPX4Deg holds promise to further enhance specific killing of cancer cells.

One of the most intriguing aspects of ferroptosis is its capacity to spread through tissues and cell populations^23–26,41^. Our optogenetics tool makes it possible to address this question in an unbiased way, meaning that, in our experiments, the non-transfected cells neighboring cells expressing Opto-GPX4Deg were not primed for ferroptosis. In this way, we could unequivocally show that ferroptotic cells can propagate cell death across the population. The accumulation of peroxidized lipids and the inhibition by ferrostatin indicated that cell death in cells not expressing Opto-GPX4Deg occurred via ferroptosis too.

Intriguingly, propagation of cell death induced by Opto-GPX4Deg occurred in a distance-dependent manner. In contrast to optogenetic induction of apoptosis, we found a clear preference for cell death induction in adjacent cells, suggesting that proximity was a key parameter in ferroptosis spread. Furthermore, our temporally and spatially-resolved data indicated that lipid peroxidation propagated to neighboring cells before cell rupture. These findings support a model in which ferroptosis can propagate to neighboring cells in direct vicinity to the dying, ferroptotic cell.

Since ferroptosis has been associated with several diseases^9–13,42,43^, our findings might have implications of medical interest. Concretely, ferroptosis propagation has been proposed to play a role in the formation of necrotic tissue in intestinal epithelium _44_, heart tissue^45–47^, excitotoxicity in brain^48^ and renal tubula^23^, which underpins the patho-physiological relevance of this cell-contact dependent process^7,23,25^. By shedding new light on the molecular mechanism of ferroptosis propagation, our results may guide new therapeutic strategies against ferroptosis-associated diseases. Furthermore, the Opto-GPX4Deg tool presented here provides new opportunities to address the physiological relevance of ferroptosis propagation *in vivo* and to deepen our understanding how ferroptosis modulators can be exploited for future treatments.

In summary, we report here a new optogenetic tool for light-controlled induction of ferroptosis, which allows initiation of this form of cell death in selected cells with unprecedented spatial and temporal resolution. Using this tool, we demonstrate that lipid peroxidation and ferroptotic cell death can propagate to neighboring cells in close proximity, which might open new possibilities for targeting ferroptosis for therapy.

### Materials and methods Plasmids

Plasmids were cloned via restriction digest cloning as follows: The pMTG02-Amp-FRT plasmid (Addgene #128271) was used as template for amplifying the LOVpep (without the RRRG degron) and LOVpepdegron fragments, which was cloned into the pcDNA™3.1 (+) backbone from Invitrogen using the ApaI and XhoI restrictions sites. Subsequently, the cDNA of GPX4 was amplified from the GPX4 plasmid (Addgene #38797) and inserted in the before generated pcDNA3.1_LOVpepdegron and pcDNA3.1_LOVpep plasmids using the EcoRI and NotI restriction sites. The cDNA of the EGFP was amplified from the pGFP-Cytochrome C (Addgene #41181) and inserted in the pcDNA3.1_GPX4_LOVpepdegron, pcDNA3.1_GPX4_LOVpep and the pcDNA3.1_LOVpepdegron to generate in the paper used GFP_GPX4_LOVpepdegron, GFP_GPX4_LOVpep and GFP_LOVpepdegron plasmids. To generate the BFP_GPX4_LOVpepdegron plasmid the mTagBFP2-TOMM20-N-10 (Plasmid #55328) was used as a template to amplify BFP2 and inserted into the GFP_GPX4_LOVpepdegron using the HindIII and EcoRI restriction sites. The GFP_Cry2 plasmid was generated by amplifying Cry2 from the Cry2(1-531)-mCh-BAXS184E plasmid (see below), which was then inserted into the GFP_LOVpepdegron using the NotI and ApaI restriction sites. The following plasmids were obtained from Addgene: Cry2(1-531)-mCh-BAXS184E (#117238), Tom20-CIB-GFP (#117242), pcDNA-NLS-PhoCl-mCherry (#87691). The insert sequences were validated by Sanger sequencing.

### Cell culture

All cell lines were cultured at 37°C in a humidified atmosphere containing 5% CO_2_. HEK293 cells (kindly provided by Frank Essmann, Univ. of Tübingen), HeLa (kindly provided by Andreas Villunger, Med. Univ. of Innsbruck) cells and the tamoxifen inducible GPX4 KO Pfa1 cells (kindly provided by Marcus Conrad, Univ. of Munich) were cultured in low-glucose Dulbecco’s modified Eagle’s medium (DMEM) (Sigma-Aldrich), and supplemented with 10% fetal bovine serum (FBS), 1% penicillin– streptomycin (P/S) (Thermo Fisher Scientific). RP285.5 murine GPX4 KO SCLC cells (kindly provided by Silvia von Karstedt, Univ. of Cologne), CT26 and B16-F19 (both kindly provided by the Pasparakis lab) were cultured in RPMI-1640 (Sigma-Aldrich), and supplemented with 10% fetal bovine serum (FBS), 1% penicillin–streptomycin (P/S) (Thermo Fisher Scientific). HeLa cells do not express E-Cadherin (https://www.proteinatlas.org/ENSG00000039068-CDH1/cell+line).

### Cell transfection

For transfections, Polyethylenimine (PEI; Polyscience, Warrington, PA) with a concentration of 1 mg/mL was used in a 5:1 ratio (PEI : DNA) for the confocal experiments and in a 3:1 for the high-throughput optogenetic experiments. In the following, the transfection procedure is described for the 8-well chambers: 25 µl Opti-MEM (Thermo Fisher) per well was mixed with 1.2 µl PEI and incubated for 2 min at room temperature. 240 ng plasmid was mixed with 25 µl Opti-MEM and subsequently added to the PEI/Opti-MEM mixture. Then the mixture was resuspended and incubated for 20 min at room temperature. Then the media from the 8-well chambers was aspirated and replaced with 200 µl DMEM media without phenol red, supplemented with 10% fetal bovine serum (FBS), 1% penicillin–streptomycin (P/S) (Thermo Fisher). Afterwards the transfection mix was added dropwise to the wells. For transfecting cells e.g. in the 6-well plate or 96 well-plate format, the above mentioned volumes of reagents were proportionally adapted and in those formats the cell culture media was not replaced on the next day, but after the transfected fresh media was supplemented with respective drugs or/and cell death markers and added to the wells.

### Confocal live cell imaging

According to workflow in **Fig. 2A**, 5.000 cells were seeded into removable 8-well chambers (Cat.No: 80841 ibidi) in combination with a glass cover slip suitable for confocal microscopy (VWR). All confocal experiments were performed using a Confocal Laser Scanning Microscope LSM 980 with Airyscan 2 and multiplex, GaAsP (4x), PMT (2x), T-PMT, external BiG.2 Typ B 980 detector (Carl Zeiss Microscopy). (Carl Zeiss Microscopy). For imaging a 40x/1,2 W C-Apochromat, Diode lasers with the wavelengths 405nm, 455nm, 488nm and 561nm (all 30 mW), GaAsP and Transmitted light (T-PMT) detectors as well as main LSM beamsplitters (MBS 405 and MBS 488/561/633) were used. The experiments were performed at 37°C and 5% CO_2_ using an integrated incubation module (Carl Zeiss Microscopy). The Image analysis software ZEN (Carl Zeiss Microscopy) was used. For acquiring confocal images, the following settings were used: Sequential acquisition for the different channels were used. The field of view was set to 512×512 pixels. For the acquisition the speed was set to max. using line-wise bidirectional scanning, whereby 4x averaging in the Repeat per Line Mode was performed. For acquiring the GFP, mCherry and BFP2 fluorescent signal the following laser setting were used: 6.5 µW for the 488 nm laser (GFP), 79 µW for the 561 nm laser (mCherry) and 13,8 µW for 445 nm laser (BFP2). Brightfield images were acquired using the 561 nm laser with 79 µW and a transmitted light (T-PMT) detector.

### Optogenetic activation in confocal microscopy

ROIs were drawn covering the whole cell area of cells expressing optogenetics tools, as well as non-transfected controls, using the ROI bleaching function of the microscope software. To control for unwanted activation, not all cells expressing the optogenetics tools within the field of view were activated. For activating the optogenetics tools, the 405 nm laser was used at 456 µW intensity according to the workflow shown in **Fig. 2A**, by setting up a time series acquiring one pre-activation image, before activating selected ROIs with 100 iterations every 7 images or with the indicated activation rounds of the respective experiment. For Bodipy quantification, confocal images were acquired 5- and 65 min upon 3 activation rounds of the optogenetic tools. For the Ca^2+^ experiments, images were acquired every 45 s upon 4 activation rounds of the optogenetic tools. Experiments were only considered for analysis if all controls worked, using blebbing of cells as proxy for cell death induction throughout the experiments. 1) Illumination of non-transfected cells to rule out phototoxicity of the illumination conditions. 2) Non-illumination of transfected cells to check for potential self-activation or construct toxicity. 3) Non-illumination of non-transfected cells to check for phototoxicity of the live cell imaging time series acquisition (background cell death). If one of these controls failed, meaning that any of the controls cells starts blebbing (indicative for cell death induction) during an experiment, the experiment was excluded from further analysis.

### High throughput optogenetic experiments

Experiments were performed according to the workflow shown in **Fig. 1B**, unless otherwise indicated. Where indicated, AnnexinV was added before a pre-activation image was captured for assessing cell death.^49^ For activation of the optogenetic tool, an optoPlate-96 was used.^29^ For programming the optoPlate-96, optoConfig-96 was used.^30^ The following illumination at 465 nm was used: 5 mW/cm^2^ 100%, 4 mW/cm^2^ for 80%, 3 mW/cm^2^ for 60%, 2 mW/cm^2^ for 40% and 1 mW/cm^2^ for 20% LED intensity. _29,30_ Kinetics of cell death and C11-Bodipy experiments were assessed using a IncuCyte S3 bioimaging platform (Essen). The pre-activation and post-activation images were acquired with the following settings: Per well, four images were captured with 400 ms exposure for the green and red channel. For assessing Bodipy 581/591, an ImageXpress Micro 4 (MD)was used. A 10x Plan Fluor 0.3 NA objective, an Andor Cycla 5.5 (CMOS) camera and the following channels were used: Tl-20 (2 ms), DAPI (50ms), FITC (200 ms) and TRITC (100 ms) with autocorrection. The experiments were performed at 37°C and 5% CO_2_ using the provided incubation chamber. For the experiment µ-Plate 24 Black ID 14 mm plates (ibidi) were used.

### Drug treatments

RSL3, ferrostatin, and Nec-2 were purchased from Biomol (Germany). zVAD was obtained from APEXBIO (Houston, TX, USA) and C11-BODIPY 581/591 as well as Flou-4-M from Thermo Fisher. Tamoxifen was bought form Sigma-Aldrich. Draq7 was obtained from Invitrogen, whereas AnnxinV (PE) was purchased from Thermo Fisher.

### Image analysis

For assessing the time till blebbing in the confocal experiments the activation time series and post activation time series were imported to Fiji and concentrated into one time series. Afterwards the channels were split to adjust Brightness and Contrast in all channels. After adjusting the channels were merged again to manually assess the time till blebbing of illuminated transfected cells, of illuminated non-transfected cells, non-illuminated of transfected cells and non-illuminated non-transfected cells.

To determine the percentage of either transfected, non-transfected or oxidized C-11 Bodipy positive cells of the high-throughput optogenetics experiments, a custom made, AI-based software was used. Thereby, the total number of transfected, non-transfected, as well as cell death-marker positive cells were calculated and normalized taking the absolute number of transfected/non-transfected cells as 100% divided by the number of double positive cells. For assessing the position and fluorescence level of individual cells, image segmentation was performed using our custom-made software. For assessing the distance between cells, the coordinates of individual cells were used by calculating the individual vector lengths.

### Lipid peroxidation measurements

Cells were transfected with Opto-GPX4Deg and incubated for 16h. On the next day the cells were stained with 1 µM Bodipy for one hour and treated with 10 µM zVAD. Then Opto-GPX4Deg expressing cells and non-transfected controls were activated as described in **(A)** for 3 times and the fluorescence level of Bodipy for individual cells (red and green) were assessed 5- and 65-min post-activation and corrected for the cell size and local background substraction was performed. The oxidation ratio of C11-Bodipy was calculated as an indicator of lipid peroxidation according to the following formula: oxidation ratio = red + green fluorescence/green fluorescence. Thereby the red fluorescence corresponds to the non-oxidized fraction of the probe and was estimated based on the fluorescence intensity per pixel from the red image and the green fluorescence corresponds to the oxidized fraction and was calculated based on the fluorescence intensity per pixel from the green channel fluorescence images. Those values were estimated based on the fluorescence intensity per pixel from the red and green channels of the fluorescence images using Fiji or our custom-made software.

### Immunoblotting

Cells were lysed in RIPA buffer supplemented with protease inhibitor (Roche). To load equal amounts of proteins on a 12 % SDS-PAGE, a Bradford assay was performed. The transfer onto a nitrocellulose membrane (Merck Millipore) was performed using a the Turboblot (BioRad) or a wet transfer system (BioRad). The membrane was blocked with 5 % milk in PBS-T (0.1 % Tween) for 1 hour at room temperature and incubated over night with the respective primary antibodies: 1:1000 ß-tubulin G8 Sc55529 (Santa Cruz), 1:1000, Anti-Glutathione Peroxidase 4 (ab125066) (Abcam), 1:1000 Anti-GFP antibody (ab290)(Abcam). The membrane was then washed five times for 5 min in PBS-T at room temperature and incubated with the secondary antibody at room temperature for one hour in the dark. The following secondary antibodies were used: 1:5000 IRDye 800CW (LI-COR) and 1:10000 IRDye 680RD (LI-COR). The membrane was washed five times for 5 min with PBS-T and the fluorescence signals were captured by an Odyssey DLx (LI-COR). Western Blots were quantified using the Image Studio Lite Software.

### Quantification of oxidized glycerophospholipids

Levels of oxidized phosphatidylcholine (PC) and phosphatidylethanolamine (PE) species were determined by Liquid Chromatography coupled to Electrospray Ionization Tandem Mass Spectrometry (LC-ESI-MS/MS) using a procedure previously described in^50^ with several modifications:

1 – 2 million cells were resuspended in 300 µl of an ice-cold solution of 100 µM diethylenetriaminepentaacetic acid (DTPA) in PBS. An aliquot of the cell suspension was used for the determination of the protein content using bicinchoninic acid. To 100 µl of the cell suspension, 2.4 ml of an ice-cold solution of 1.5 mg/ml triphenylphosphine and 0.005 % butylated hydroxytoluene in methanol were added. The mixture was incubated for 20 min in a shaking bath at room temperature. Afterwards, 1 ml of the 100 µM DTPA in PBS solution, 1.25 ml of chloroform and internal standards (10 pmol 1,2-dimyristoyl-sn-glycero-3-phosphocholine (DMPC) and 10 pmol 1,2-dimyristoyl-sn-glycero-3-phosphoethanolamine (DMPE)) were added. The samples were vortexed for 1 min and incubated at -20 °C for 15 min. After adding 1.25 ml of chloroform and 1.25 ml of water, the mixture was vortexed vigorously for 30 sec and then centrifuged (4,000 × g, 5 min, 4 °C) to separate layers. The lower (organic) phase was transferred to a new tube and dried under a stream of nitrogen. The residues were resolved in 150 µl of methanol and transferred to autoinjector vials.

LC-MS/MS analysis was performed as previously described.^50,51^ The LC chromatogram peaks of oxidized PC and PE species and the internal standards were integrated using the MultiQuant 3.0.2 software (SCIEX). Oxidized PC and PE species were quantified by normalizing their peak areas to those of the internal standards. The normalized peak areas were then normalized to the protein content of the cell suspension.

### Quantification of fatty acids

To 50 µl of the cell suspension mentioned above, 50 µl of water, 500 µl of methanol, 250 µl of chloroform, and internal standard (1 µg palmitic-d31 acid) were added. The mixture was sonicated for 5 min, and lipids were extracted in a shaking bath at 48 °C for 1 h. Glycerolipids were degraded by alkaline hydrolysis adding 75 µl of 1 M potassium hydroxide in methanol. After 5 min of sonication, the extract was incubated for 1.5 h at 37 °C, and then neutralized with 6 µl of glacial acetic acid. 2 ml of chloroform and 4 ml of water were added to the extract which was vortexed vigorously for 30 sec and then centrifuged (4,000 × g, 5 min, 4 °C) to separate layers. The lower (organic) phase was transferred to a new tube, and the upper phase extracted with additional 2 ml of chloroform. The combined organic phases were dried under a stream of nitrogen. The residues were resolved in 300 µl of acetonitrile/water 2:1 (v/v) and sonicated for 5 min. After centrifugation (12,000 × g, 2 min, 4 °C), 40 µl of the clear supernatants were transferred to autoinjector vials.

Fatty acid levels were determined by LC-ESI-MS/MS using a procedure previously described in^52^with a few modifications: 10 µl of sample were loaded onto a Core-Shell Kinetex Biphenyl column (100 mm × 3.0 mm ID, 2.6 µm particle size, 100 Å pore size, Phenomenex), and fatty acids were detected using a QTRAP 6500 triple quadrupole/linear ion trap mass spectrometer (SCIEX). The LC (Nexera X2 UHPLC System, Shimadzu) was operated at 40 °C and at a flow rate of 0.7 ml/min with a mobile phase of 5 mM ammonium acetate and 0.012 % acetic acid in water (solvent A) and acetonitrile/isopropanol 80:20 (v/v) (solvent B). Fatty acids were eluted with the following gradient: initial, 55 % B; 4 min, 95 % B; 7 min, 95 % B; 7.1 min, 55 % B; 10 min, 55 % B.

Fatty acids were monitored in the negative ion mode using “pseudo” Multiple Reaction Monitoring (MRM) transitions.^52^ The instrument settings for nebulizer gas (Gas 1), turbogas (Gas 2), curtain gas, and collision gas were 60 psi, 90 psi, 40 psi, and medium, respectively. The interface heater was on, the Turbo V ESI source temperature was 650 °C, and the ionspray voltage was -4 kV.

The LC chromatogram peaks of the endogenous fatty acids and the internal standard palmitic-d31 acid were integrated using the MultiQuant 3.0.2 software (SCIEX). Endogenous fatty acids were quantified by normalizing their peak areas to those of the internal standard. The normalized peak areas were then normalized to the protein content of the cell suspension.

### Statistical analysis

For assessing significant differences different statistical tests were performed depending on the number of compared groups as well as on the factors considered in the experiment. For comparing two groups or conditions non-parametric t-tests were performed. When more than one t-test was performed within on graph, they were corrected for multiple comparisons using a multiple t-test. When more than two groups or conditions were analyzed, a one-way ANOVA was performed, when one factor was compared, whereas and a three-way ANOVA was performed, when three factors were compared. For correcting for multiple comparisons, the Tukey’s multiple comparison test was performed. For performing the statistical tests, Grappad Prism 7.03 was used. For assessing the clustering tendency of the data set, Hopkins statistics was performed using freely available Code on Github.^53^

## Acknowledgments

This project was funded by the Deutsche Forschungsgesellschaft (DFG, German Research foundation), SFB1403 – project no. 414786233 and partially SPP2306 – project no. GA1641/7-1 and by the Bundesministerium für Bildung und Forschung (BMBF) project. 16LW0213. We also thank the CECAD Imaging and Lipidomics/Metabolomics Facilies as well as staff members for their support.

## Author contributions

BFR performed experiments and analyzed data. MRHV built custom software and analyzed experiments. All authors contributed to experimental design and manuscript writing. AJG-S conceived the project, supervised research and wrote the manuscript together with BFR.

