## Supplementary figures 1-8 for "Ferroptosis propagates to neighboring cells via cell-cell contacts"

### Supplemtary Figures

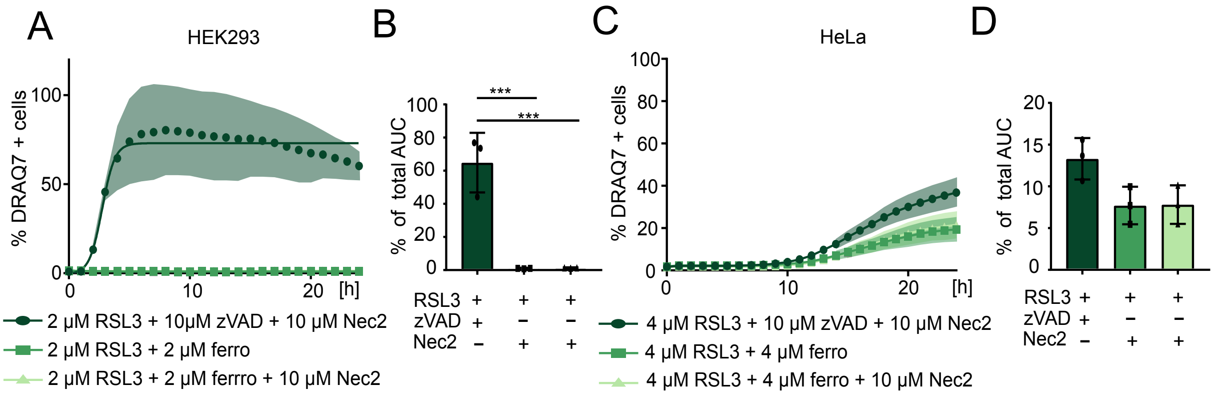


**Figure S1. (A-D) Chemical ferroptosis induction in HEK and HeLa cells**. On day 0, 10.000 HeLa or HEK293 cells/well were seeded into a 96-well plate. On the next day, HeLa or HEK cells were treated with different combination treatments: (RSL3 + zVAD + Nec2), (RSL3 + ferrostatin) or (RSL3 + ferrostatin + Nec2) at the in indicated concentrations and 1:200 Draq7 was added for cell death assessment. Cell death kinetics assessed via Draq7 staining and using a IncuCyte S3. To assess statistical differences, % total area under the curve of different treatments was calculated and subsequently a one-way ANOVA was performed. Asterisks indicate significant differences: *p < 0.05, **p < 0.01, ***p < 0.001, ****p<0.0001. n = 3 for all experiments. Values are displayed as mean ± SD.

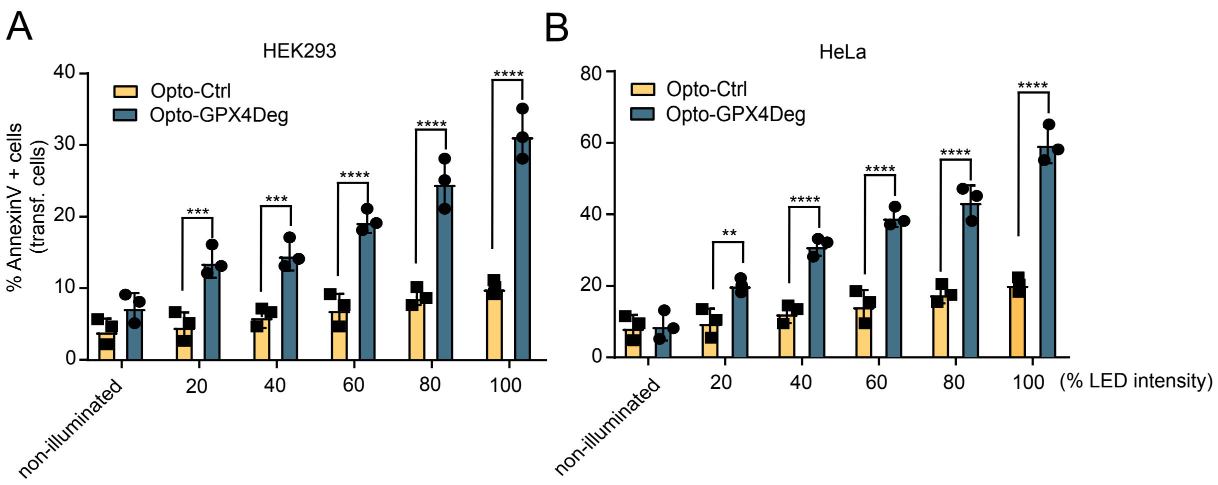


**Figure S2. (A and B) Increased illumination intensity leads to proportionally increased cell death induction in cells expressing Opto-GPX4Deg compared to Opto-Ctrl.** Experiments performed according to **Fig. 1B** except that a gradient of 465 nm LED intensities was used. Cell death assessed by AnnexinV staining. % AnnexinV positive cells normalized to GFP positive cells 10h post activation. For assessing statistical differences, a multiple t-test using the Holm-Sidak method was performed. Asterisks indicate significant differences: *p < 0.05, **p < 0.01, ***p < 0.001, ****p<0.0001. n = 3 for all experiments. Values are displayed as mean ± SD.

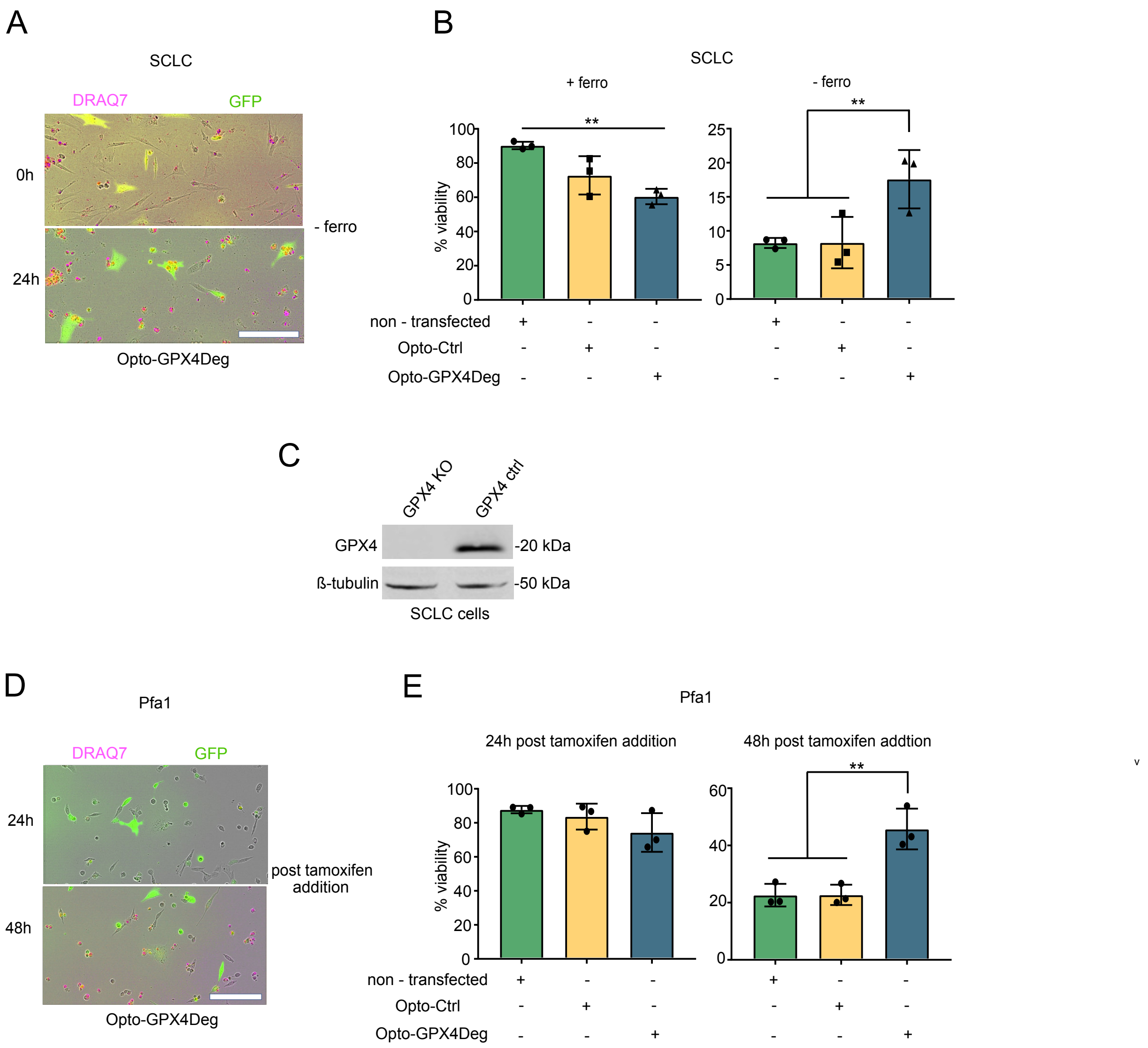


**Figure S3. Opto-GPX4Deg retains partial GPX4 activity. (A)** To test the functionality of the GPX4 within the optogenetic fusion protein, 5.000 RP285.5 murine GPX4 KO SCLC cells/well were seeded into a 96-well plate. On the next day, the cells were either transfected or not with Opto-GPX4Deg or Opto-Ctrl and cultured with or without 5 µM ferrostatin supplementation for 24h. On the next day, 1:200 Draq7 was added to the cells and cell death was assessed. We quantified the % viable, non-transfected cells expressing Opto-GPX4Deg cultured with or without 5 µM ferrrostatin and % viable, non-transfected cells expressing Opto-Ctrl. For assessing statistical differences, a one-way ANOVA was performed and corrected for multiple comparisons using Tukey's multiple comparison test. Asterisks indicate significant differences: *$p < 0.05$, **$p < 0.01$, ***$p < 0.001$, ****$p<0.0001$. n = 3 for all experiments. Values are displayed as mean ± SD. **(B)** Representative images showing increased viability in Opto-GPX4Deg expressing cells compared to non-transfected cells cultured in ferrostatin deprived medium. **(C)** Representative WB showing that the SCLC GPX4 KO cells are GPX4 deficient. **(D and E)** As alternative approach for testing the GPX4 function within the optogenetic fusion protein, 2.500 Pfa1 cells (a tamoxifen inducible GPX4 KO cell line) were seeded into a 96 well-plate and transfected on the next day with Opto-GPX4Deg in presence of 1 μM tamoxifen. 1:200 Draq7 was used to calculate cell death. **(D)** Representative images showing increased viability in Opto-GPX4Deg expressing cells compared to non-transfected cells upon 48h tamoxifen induction. **(E)** Statistical analysis by one-way ANOVA corrected for multiple comparisons using Tukey's multiple comparison test. Asterisks indicate significant differences: *$p < 0.05$, **$p < 0.01$, ***$p < 0.001$, ****$p<0.0001$. n = 3 for all experiment. Values are displayed as mean ± SD.

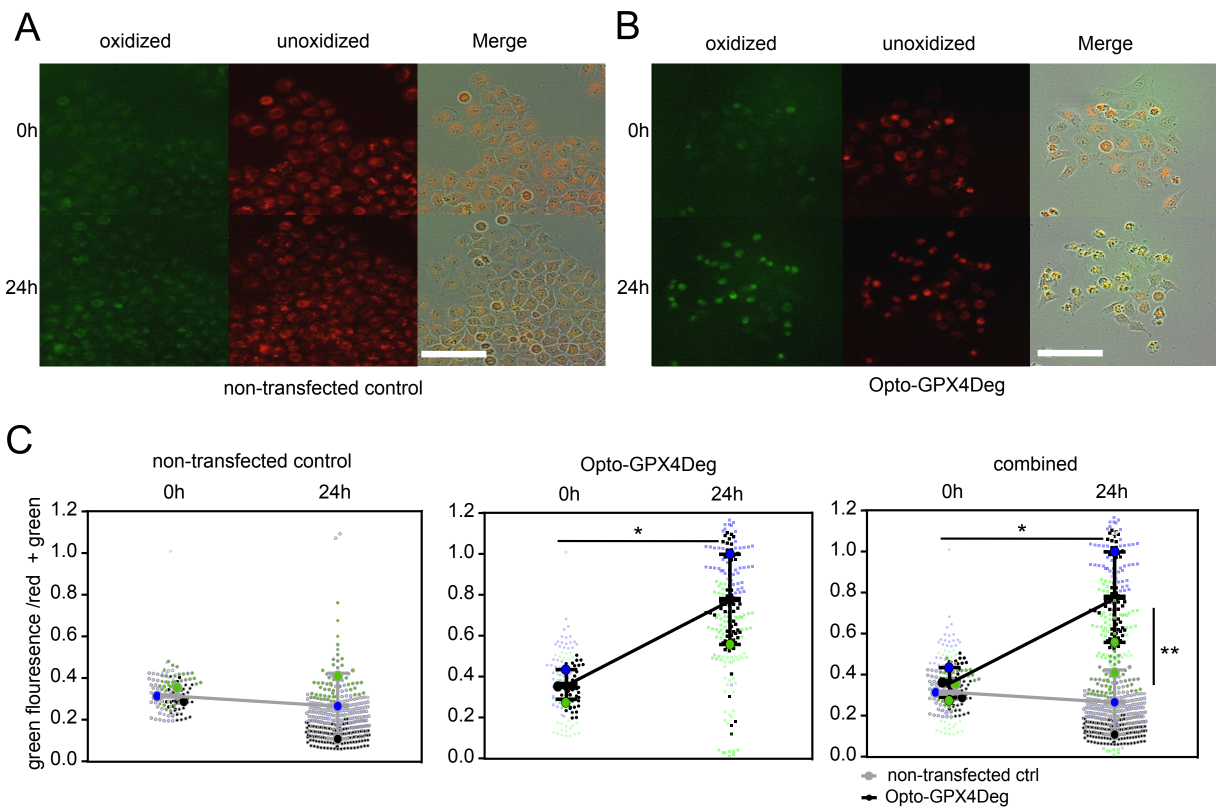


**Figure S4. High-throughput optogentic ferroptotic induction results in elevated levels of lipid peroxidation. (A - C)** C11-Bodipy quantification. On the first day, 100.000 cells were seeded into 12-well plates and transfected or not on the next day with Opto-GPX4Deg, in this case BFP2_GPX4_LOVpepdegron, and incubated for 16h at 37°C. On the subsequent day, HeLa cells were stained with 1 µM C11-Bodipy for 1h and then activated for 30 min with 100% 465 nm LED intensity using the optoPlate-96. For normalization, the green (oxidized) C11-Bodipy signal was quantified, divided by the sum of red and green C11-Bodipy signal (total). The big dots represent the mean of three independent experiments, the small dots represent single cells in one individual experiment. Statistica analysis by one-way ANOVA corrected for multiple comparisons using Tukey's multiple comparison test. Asterisks indicate significant differences: *$p < 0.05$, **$p < 0.01$, ***$p < 0.001$, ****$p<0.0001$. Values are displayed as mean ± SD.

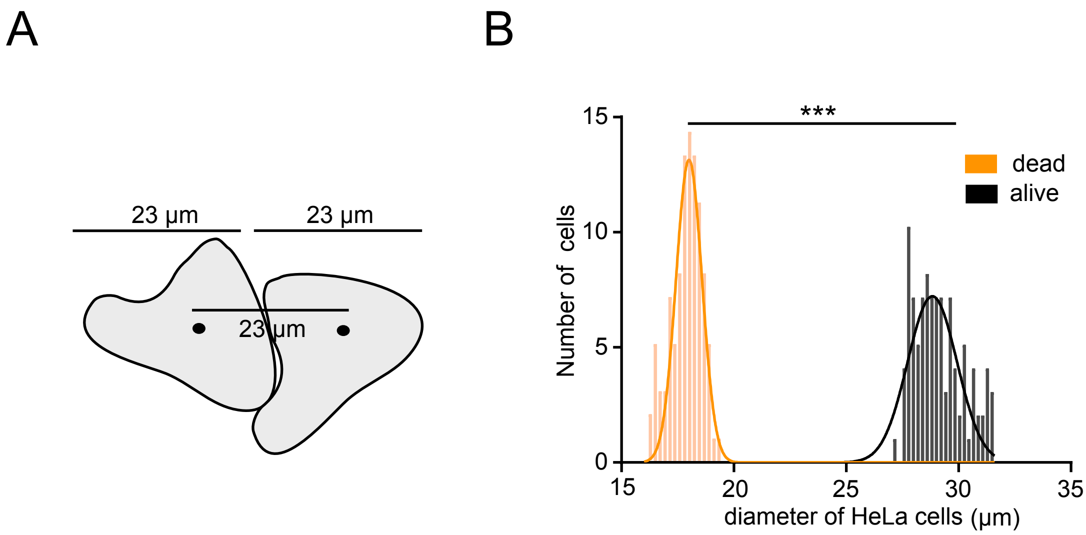


**Figure S5: HeLa cell have an approximate average size of 23 µm (A)** Scheme indicating the average diameter of HeLa cells as well as the average distance between adjacent HeLa cells. **(B)** Quantification of the cell diameter distribution of living and dead HeLa cells derived from the experiments in **Fig.5 A-C.** For assessing statistical differences, a parametric t-test was performed. Asterisks indicate significant differences: *$p < 0.05$, **$p < 0.01$, ***$p < 0.001$, ****$p<0.0001$. n = 3 for all experiments. Values are displayed as mean ± SD.

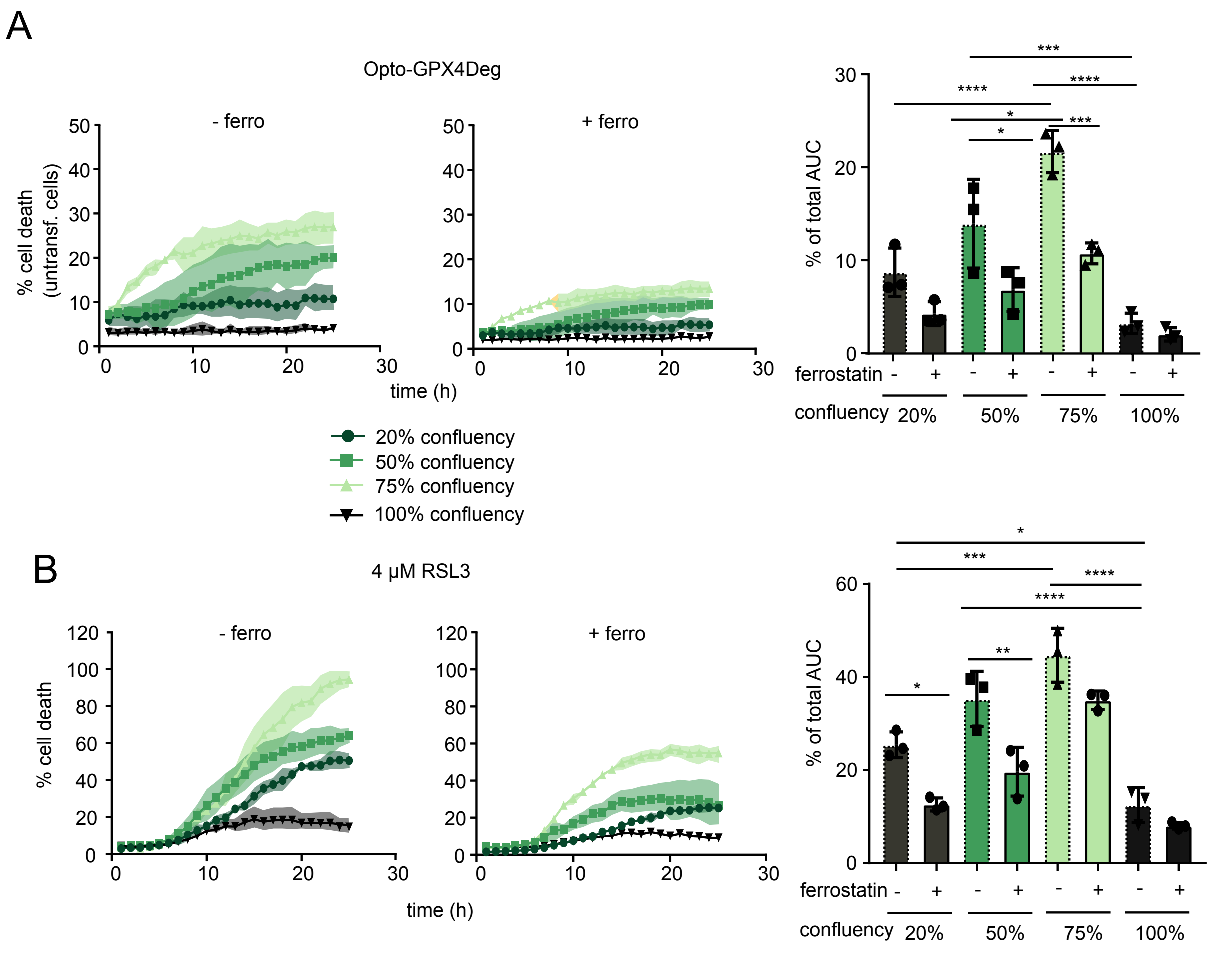


### Figure S6: Increased cell confluency renders HeLa cells more sensitive to ferroptosis

**(A)** Optogenetic experiments were performed as described in (**Fig. 1B)** with the difference that here either 2.5, 5, 10 or 15k HeLa cells were seeded into 96-well plate and subsequently incubated for 1 days before being transfected with Opto-GPX4Deg for 16h. On the next day the cell lines were treated or not treated with 5 µM ferrostatin and 1:200 Draq7 was added for cell death assessment. Cell death kinetics was assessed via Draq7 staining and using a IncuCyte S3. **(B)** On day 0 2.5, 5, 10 or 15k HeLa cells per well were seeded into a 96-well plate. On the next day the cells were treated with different combination treatments: RSL3, RSL3 + ferrostatin or DMSO in the in indicated concentrations and 1:200 Draq7 was added for cell death assessment. To assess statistical differences the % of total area under the curve of the different treatments was calculated and subsequently a one-way ANOVA was performed and corrected for multiple comparisons using Tukey's multiple comparison test. Asterisks indicate significant differences: *$p < 0.05$, **$p < 0.01$, ***$p < 0.001$, ****$p<0.0001$. n = 3 for all experiments. Values are displayed as mean ± SD.

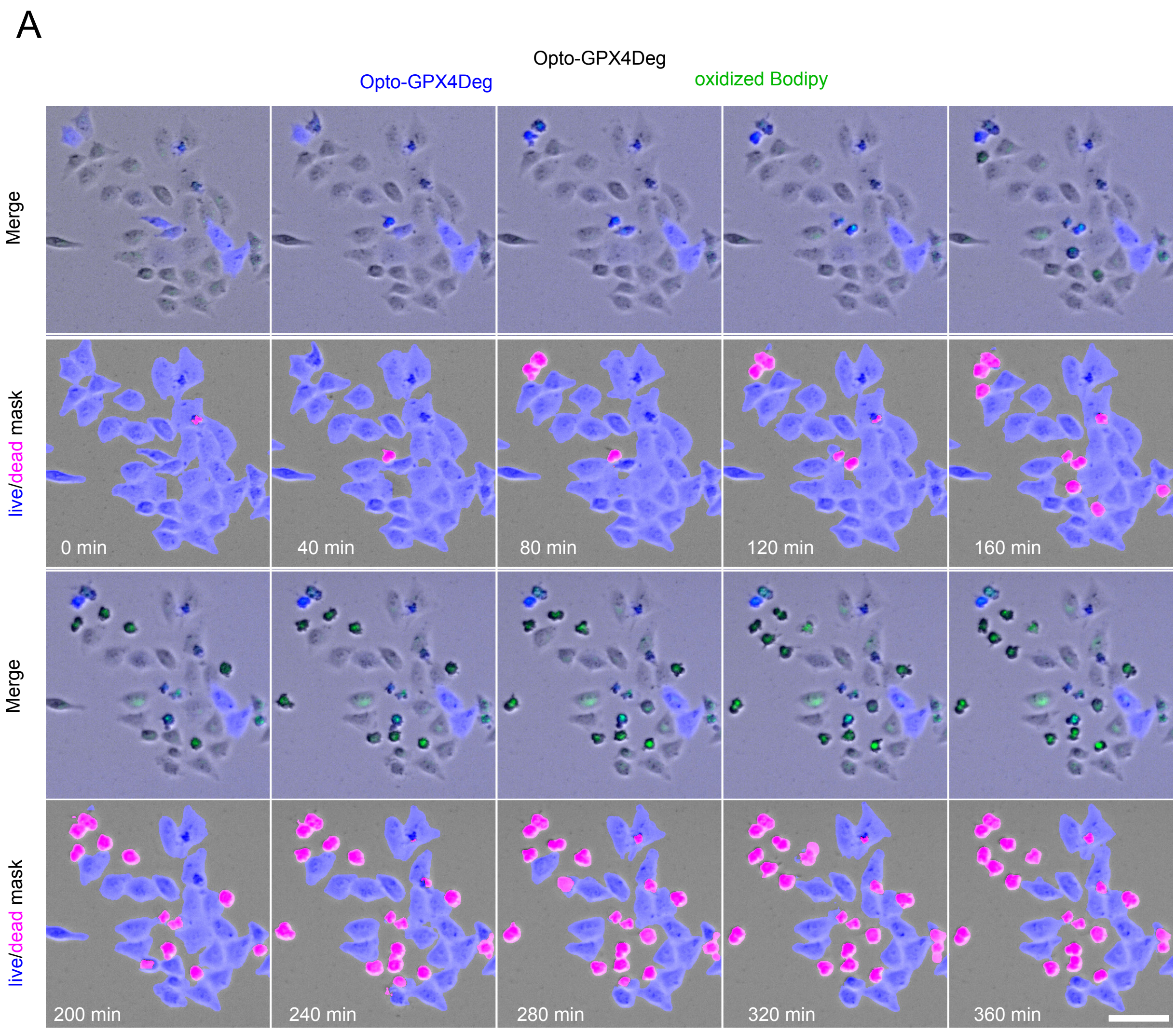


**Figure S7: AI derived live/dead mask enables the assessment of cell death kinetics.** Data is derived from the experiment performed in **Fig.6** Representative Bodipy/cell death kinetic time series of the Opto-GPX4Deg tool expressing bulk. Blue signal indicates for expression of the optogenetic tool and the green signal for the presence of lipid peroxidation (upper panel). (Lower panel) Cell death output overlay of the AI based software. Living cells are indicated in blue and dead cells in pink. Scale bars, 100 μm.

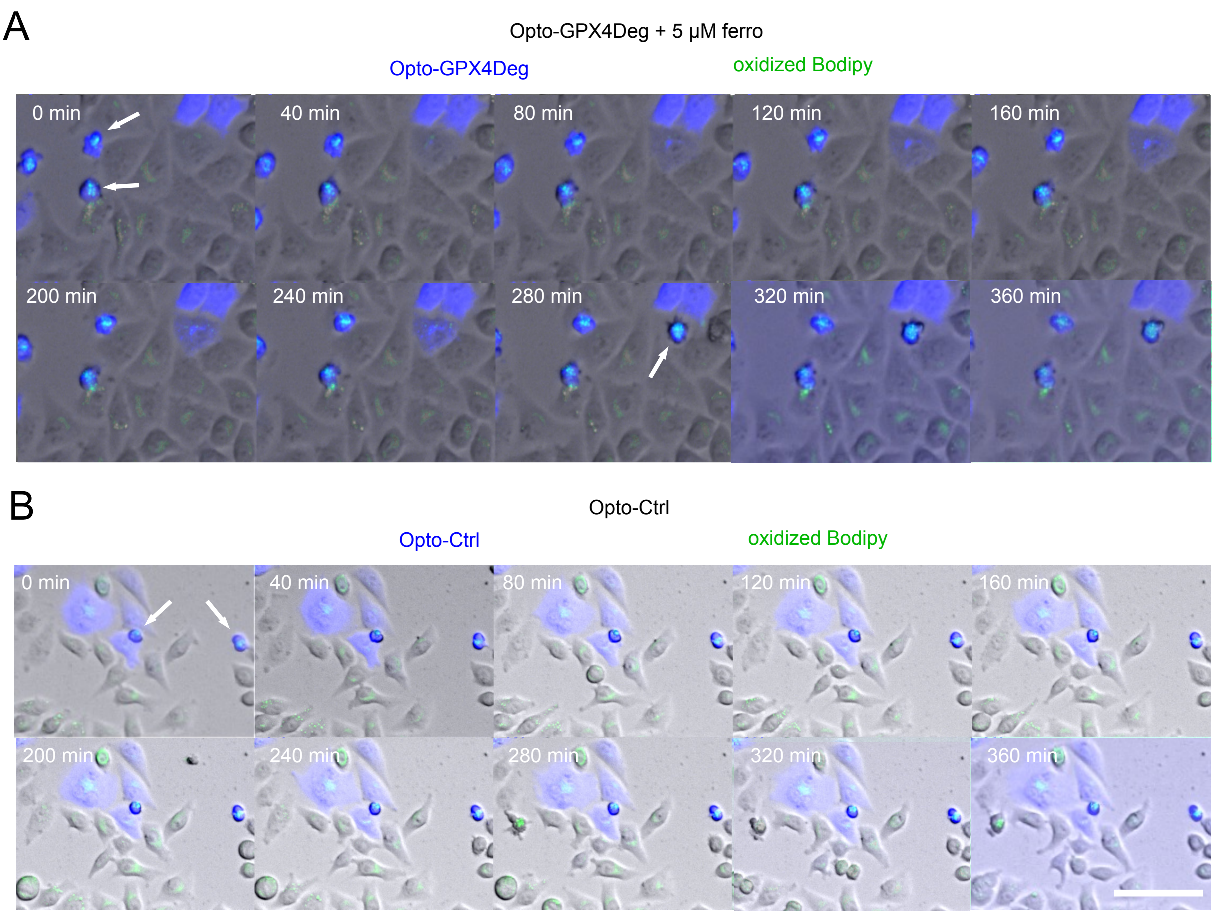


**Figure S8: Ferrostatin blocks cell death propagation to neighboring cells**

Data is derived from the experiment performed in **Fig.6** Representative C11-Bodipy/cell death kinetic time series of the indicated condition. Blue signal indicates for expression of the optogenetic tool and the green signal for the presence of lipid peroxidation. Green arrows indicate dead, lipid peroxide positive cells, whereas black arrows indicate dead lipid peroxide negative cells. Scale bars, 100 µm.
